# *In Vivo* Multi-Day Calcium Imaging of CA1 Hippocampus in Freely Moving Rats Reveals a High Preponderance of Place Cells with Consistent Place Fields

**DOI:** 10.1101/2021.08.17.456533

**Authors:** Hannah S Wirtshafter, John F Disterhoft

**Affiliations:** Department of Neuroscience, Northwestern University Feinberg School of Medicine, Chicago, IL, USA

## Abstract

Calcium imaging using GCaMP indicators and miniature microscopes has been used to image cellular populations during long timescales and in different task phases, as well as to determine neuronal circuit topology and organization. Because the hippocampus (HPC) is essential for tasks of memory, spatial navigation, and learning, calcium imaging of large populations of HPC neurons can provide new insight on cell changes over time during these tasks. All reported HPC in vivo calcium imaging experiments have been done in mouse. However, rats have many behavioral and physiological experimental advantages over mice. In this paper, we present the first (to our knowledge) in vivo calcium imaging from CA1 hippocampus in freely moving male rats. Using the UCLA Miniscope, we demonstrate that, in rat, hundreds of cells can be visualized and held across weeks. We show that calcium events in these cells are highly correlated with periods of movement, with few calcium events occurring during periods without movement. We additionally show that an extremely large percent of cells recorded during a navigational task are place cells (77.3±5.0%, surpassing the percent seen during mouse calcium imaging), and that these cells enable accurate decoding of animal position and can be held over days with consistent place fields in a consistent spatial map. A detailed protocol is included, and implications of these advancements on *in vivo* imaging and place field literature are discussed.

**Significance statement:** In vivo calcium imaging in freely moving animals allows the visualization of cellular activity across days. In this paper, we present the first in vivo Ca2+ recording from CA1 hippocampus in freely moving rats. We demonstrate that hundreds of cells can be visualized and held across weeks, and that calcium activity corresponds to periods of movement. We show that a high percentage (77.3±5.0%) of imaged cells are place cells, and that these place cells enable accurate decoding and can be held stably over days with little change in field location. Because the hippocampus is essential for many tasks involving memory, navigation, and learning, imaging of large populations of HPC neurons can shed new insight on cellular activity changes and organization.

## Introduction

The explosion in the use of *in vivo* calcium imaging has allowed the visualization of hundreds of identified neurons over long time periods (Grienberger and Konnerth, 2012a; Grienberger and Konnerth, 2012b; Rickgauer et al., 2014; Hamel et al., 2015; Resendez et al., 2016; Aharoni and Hoogland, 2019b; Aharoni et al., 2019; Zhou et al., 2019). Calcium (Ca2+) imaging using GCaMP calcium indicators and miniature microscopes has been used to image cellular populations during long timescales (Dombeck et al., 2010; Ziv et al., 2013; Cai et al., 2016; Sheintuch et al., 2017; Hainmueller and Bartos, 2018; Kinsky et al., 2020; Sheintuch et al., 2020) and in different task phases (Zhang and Li, 2018; Jimenez et al., 2020; Shuman et al., 2020), as well as to determine neuronal circuit topography and organization (Klaus et al., 2017; Wang et al., 2017; Guo et al., 2020; Shin et al., 2020).

The hippocampus (HPC) has been studied with cellular resolution using *in vivo* electrophysiology, but this technique does not allow the same cells to be definitively followed over days, nor does it allow the visualization of cellular organization within the structure (O’Keefe, 1976; Wilson and McNaughton, 1993; McEchron and Disterhoft, 1999; Leutgeb et al., 2004; Schimanski et al., 2013). *In vivo* calcium imaging is particularly well suited for imaging the rodent HPC, as the orientation of the horizontal cell layer permits imaging of large numbers of neurons with insertion of a 1mm diameter or smaller lens (Guo et al., 2020; Kinsky et al., 2020; Sheintuch et al., 2020; Stefanini et al., 2020). Because the hippocampus is essential for many tasks involving memory (Scoville and Milner, 1957; Squire, 1992; Eichenbaum et al., 1999; Hasselmo and McClelland, 1999; Ferbinteanu and Shapiro, 2003; Ferbinteanu et al., 2006; Smith and Mizumori, 2006; Vann, 2013; Cai et al., 2016; Josselyn and Tonegawa, 2020), spatial navigation (Foster et al., 2000; Rosenzweig et al., 2003; McNaughton et al., 2006; Foster and Knierim, 2012; Moser et al., 2017; Avigan et al., 2020), and learning (Moyer et al., 1990; Chan et al., 2001; Ito et al., 2005; Andrzejewski and Ryals, 2016), Ca2+ imaging of large populations of HPC neurons can shed new insight on cell changes and organization over time during these tasks.

To our knowledge, all reports of hippocampal *in vivo* calcium imaging have been done in mouse (although calcium imaging has been used to visualize other structures in rat (Scott et al., 2018; Anner et al., 2020; Hart et al., 2020; Pritchard et al., 2021)). Using calcium imaging in rat hippocampus requires optimizing a number of methodological parameters, including aspiration and lens implantation techniques, to account for the increased thickness of the rat alveus compared to mouse which makes focusing on the CA1 cell layer difficult (Swanson, 2004; Routh et al., 2009; Franklin and Paxinos, 2013). However, rats have many basic experimental advantages over mice: First, rats are able to learn and perform more complex tasks than mice, and employ more advanced strategies, and are thus suited for a wider variety of behavioral experiments (Whishaw, 1995; Frick et al., 2000; Whishaw et al., 2001; Cressant et al., 2007; Rosenfeld and Ferguson, 2014; Hok et al., 2016). Second, compared to mouse physiology, rat physiology for a number of conditions and disorders (including addiction, aging, and schizophrenia, among others) more closely mirrors human physiology (Iannaccone and Jacob, 2009; Vengeliene et al., 2014; Ellenbroek and Youn, 2016; Carter et al., 2020). Third, rats are less stressed by handling than mice, thus introducing less variability in experimental procedures (Buerge and Weiss, 2004).

Rats are also an advantageous organism to use for calcium imaging: Due to their larger size, rats are able to support larger implants, including a wider diameter imaging lens, larger and heavier cameras that may be available in the future, and more combinations with electrophysiology implants. This advantage will thereby enable the recording of a greater number of cells (Voigts et al., 2013; Aharoni and Hoogland, 2019b; Voigts et al., 2020). Additionally, rat hippocampal neurons are less excitable than those in mice (Routh et al., 2009; Hok et al., 2016), and rats are less prone to post-operative seizures than mice (Hunt et al., 2009, 2010; Bolkvadze and Pitkanen, 2012). Rats may therefore, compared to mice, be at a decreased risk of the seizures (and potentially to cell death) seen in mice after prolonged GCaMP expression (Tian et al., 2009b; Grienberger and Konnerth, 2012b; Resendez et al., 2016; Steinmetz et al., 2017; Yang et al., 2018).

Although *in vivo* Ca2+ imaging would appear to hold substantial promise for the study of hippocampus in freely moving rats, there is currently no direct evidence that it will work in this system. To our knowledge, there are no published studies employing *in vivo* Ca2+ imaging in rat hippocampus, likely partially due to the number of methodological changes necessary to adapt the technique from mice to rats. In this paper, we present reports of *in vivo* Ca2+ recording from CA1 hippocampus in freely moving rats. We demonstrate that hundreds of cells are reliably visualized and held across weeks using this technique, and that calcium activity corresponds to periods of movement. We show that during a navigational task involving retrieving food reward on a linear track, a uniquely high percentage (77.3±5.0%) of imaged cells are place cells (as compared to calcium imaging studies in mouse and electrophysiology studies in rat and mouse), and that these place cells enable accurate decoding and can be held stably over days with little change in field location. A detailed protocol for this technique, including notes on the numerous parameter changes needed to use Ca2+ in rats, is included in the Materials and Methods section, and implications of these advancements are discussed.

## Materials and Methods

### SUBJECT DETAILS

All procedures were performed within Northwestern Institutional Animal Care and Use Committee and NIH guidelines. Five male Fischer 344 x Brown Norway rats (275g to 325g) were sourced from National Institute on Aging colony at Charles River Laboratories, injected with AAV9-GCaMP7c, implanted with a 2mm GRIN lens and run on a linear maze (Figure 1). Animals were individually housed in an animal facility with a 12h light dark cycle.

**Figure 1:**
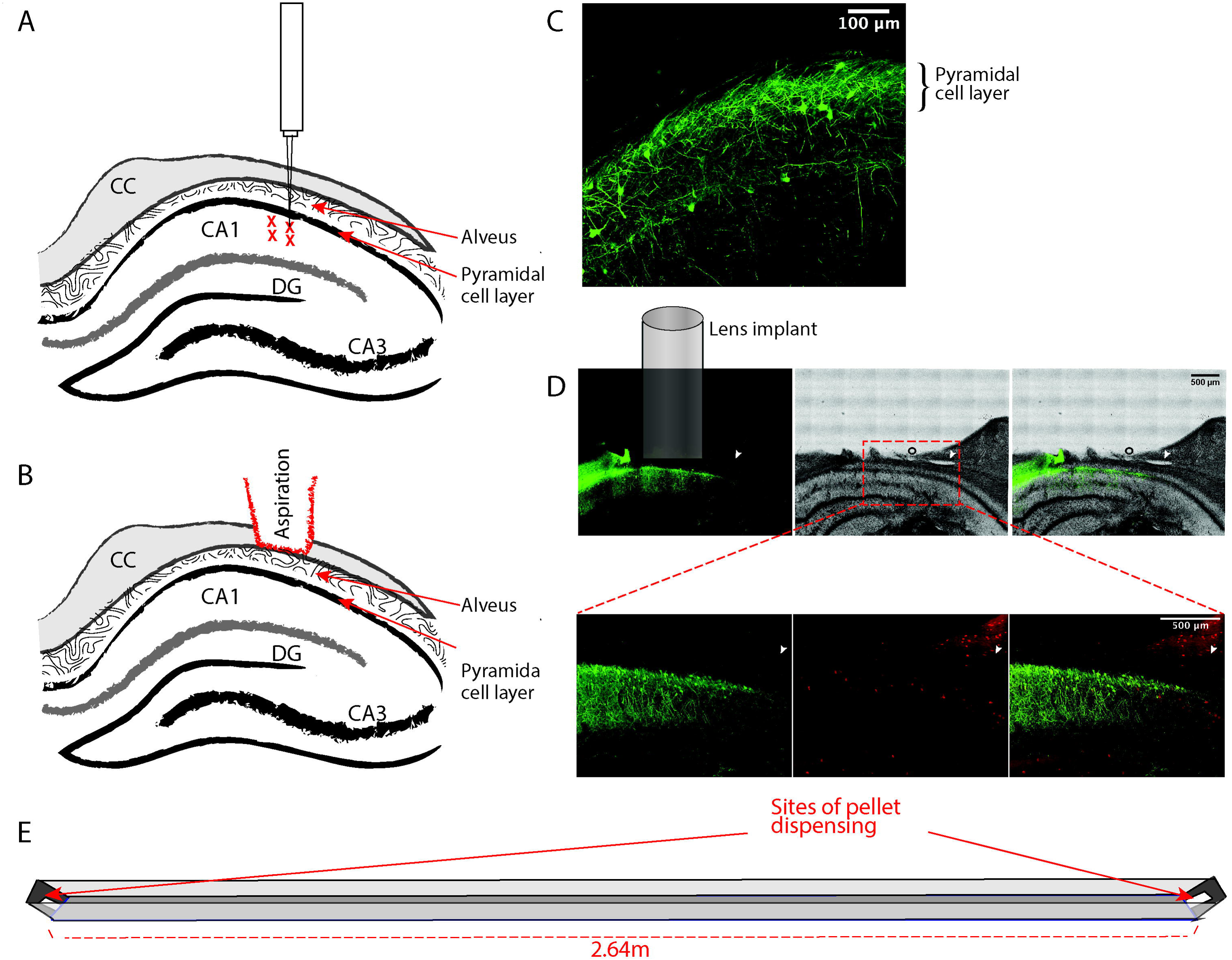
GCaMP is expressed in the CA1 cell layer of rat hippocampus. a. Diagram showing injection locations of AAV9-Syn-GCaMP7c, right below the CA1 cell layer. There are four total injection sites, with 0.6uL injected at each site. (CC = corpus callosum, DG = dentate gyrus) b. Diagram showing level of aspiration in brain tissue. The entire corpus callosum is aspirated until the vertical striations of the alveus are visible. c. GCaMP expression in the hippocampus, taken with a confocal microscope. The CA1 cell layer is labeled. d. Cellular GCaMP expression is primarily localized to principal cells in the pyramidal call layer. All images are from the same section; arrow indicates a space in the tissue that can be followed in all images. Top: a comparison of GCaMP labeling (left) with cresyl violet staining (middle) performed sequentially in the same tissue slice. An overlay of the two images (right) shows that cellular GCaMP expression is highest in the pyramidal cell layer while labeled dendrites extend ventrally into the stratum radiatum. The outlined area in the middle image roughly corresponds to the area imaged in the bottom panel. Bottom: GCaMP labeling (left) and labeling of parvalbumin+ (PV+) interneurons using immunohistochemistry (center). The overlay (right) indicates only a minority of GCaMP cells in the stratum pyramidale expressed parvalbumin. The majority of PV+ cells appear to be located slightly ventral to the cellular GCaMP labeling. e. Diagram of linear track. The animal would run back and forth between the two sides to receive a pellet reward

### METHOD DETAILS

#### GCaMP7c Injection and Lens Implantation

Rats were anesthetized with isoflurane (induction 4%, maintenance 1-2%). Following anesthesia, ear bars, and skull leveling, the skull was cleaned and a craniotomy site was marked above CA1 (stereotaxic coordinates Bregma AP -4.00mm, ML 3.00mm). The skull was scored around the craniotomy site using a razor blade to provide texture for future application of epoxy. A craniotomy was made with a 2mm trephine around the marked site and dura was removed with a bent 30 gauge needle. A Hamilton syringe (Gastight® 1700 Series Syringes, Hamilton, Model 1705 Small RN Syringe, Volume=50 µL, Point Style=3, Gauge=22s, Needle Length=50.8 mm) loaded with AAV9-GCaMP7c (obtained from Viogene Biosciences, packaged AAV9 of pGP-AAV-syn-jGCaMP7c-WPRE, titer 1.73*10^13^GC/mL) was lowered straight down to about 0.2mm past the CA1 cell layer (approximate coordinates Bregma AP -4.00mm, ML 3mm, DV 2.95mm relative to skull). 0.6uL of virus were injected over 12 minutes using an automated stereotaxic Injector. The syringe was then raised 0.2mm (Bregma AP - 4.00mm, ML 3mm, DV 2.75mm) and another 0.6uL of virus were injected over 12 minutes (Figure 1a). The syringe was then left for 10 minutes before slowly raising to the skull.

We used a computerized stereotaxic injection system that allowed us to specify the coordinates of the injection. Due to the imprecision of craniotomies and imperfectly level skulls, often the initial injection at the specified sub-cortical coordinates did not fall in the center of the craniotomy. To increase and maximize the spread of viral injection, we completed a second injection. The process of lowering the syringe and doing two injections was repeated so that the two syringe “lowering points” were evenly spread throughout the craniotomy site. There were thus a total of four injections of 0.6uL each (2.4uL total): two injections per each brain entrance site, each at different depths (Figure 1a).

Following injection, four skull screws were screwed into the skull: one rostral to bregma above the same hemisphere as the craniotomy, two between bregma and lambda on the hemisphere opposite the craniotomy, and one behind lambda on the hemisphere with the craniotomy. Under a dissecting microscope, tissue from the craniotomy was then aspirated using a vacuum pump and 25 gauge needle that had been previously blunted using a Dremel drill. Tissue was quickly aspirated until the start of horizontal striations could be seen in the craniotomy hole. Care was then taken to “level” the hole and make sure aspiration was at an even depth. The horizontal striations of the corpus callosum were then carefully and slowly aspirated until the vertical striations of the alveus were visible in the entirety of the craniotomy hole (Figure 1b). If the aspiration hole was bleeding, a small twisted piece of gel foam was inserted into the hole for approximately five minutes and then removed.

> Notes differentiating this method from mouse Ca2+ imaging: We found the best AAV for expression in rats was AAV9. Because of rat’s lower seizure threshold and, anecdotally noted, lower GCaMP expression than in mice, four injection sites appear both necessary and sufficient for ideal GCaMP expression. The injection volumes are much larger than those used in mice (Ziv et al., 2013; Cai et al., 2016), which is made possible due to the larger hippocampal area and the technique of injecting below the CA1 cell layer, instead of on it. Unlike in mice, where just the cortex or most dorsal part of the corpus callosum is aspirated (Cai et al., 2016; Schoenfeld et al., 2021), it is important that the entire corpus callosum be completely aspirated in order to get very close to the alveus.

A 2mm GRIN lens (obtained from Go!Foton, CLH lens, 2.00mm diameter, 0.448 pitch, working distance 0.30mm, 550nm wavelength) was then held on the end of a pipette using suction from the vacuum pump. The pipette was placed in the stereotaxic instrument which was zeroed with the bottom of the lens on bregma. The lens was then moved to the center of the craniotomy and lowered to a depth of approximately 3mm from skull surface. The lens was then fixed in place with UV curing epoxy and the vacuum pump was turned off and the pipette holding the lens removed. The epoxied lens was then secured to the skull screws using dental acrylic. Animals were then given buprenorphine (0.05mg/kg) and 20mL saline, taken off anesthesia, and allowed to recover in a clean cage placed upon a heat pad.

> Notes differentiating this method from mouse Ca2+ imaging: A much larger lens (2mm diameter as opposed to 0.5mm or 1.0mm diameter) lens can be used in rats due to their larger size. Of high importance: in order to be as close to the CA1 layer as possible, the lens is inserted further down (about .5mm) lower than the cell layer. This serves, presumably, to temporarily compress the alveus to make sure the alveus is pressed against the grin lens; this allows for in focus imaging of the CA1. We saw no permanent compression of the CA1 cell layer (Figure 1C) or any behavioral deficits that are associated with HPC compression (Shim et al., 2003; Chen et al., 2017). We had zero visualization of cells unless the lens was deeply inserted. Additionally, unlike in mouse implantation surgeries, we did not have success using meloxicam after surgery as it appeared to cause excessive bleeding around the lens.

Four to six weeks after surgery, animals were again anesthetized with isoflurane and checked for cells expressing GCaMP. If expression was seen, the animal’s lens was base-plated. The baseplate was attached using UV-curing epoxy and dental acrylic.

> Notes differentiating this method from mouse Ca2+ imaging: During base plating, it gets progressively harder to see cells the longer the animal is under anesthesia. Additionally, while a few cells will appear to be bright circles in the captured image, cells do not appear to be flashing at all while the rat is under isoflurane, even after a tail pinch or loud noise (techniques often used to incite a calcium burst during mouse base plating). We recommend focusing on the bright, not flashing cells while base plating. The entire field of view should become progressively brighter as the animal wakes up. We recommend not imaging for experiments within 24 hours of isoflurane exposure and base plating.

#### Behavioral Training

Prior to training, animals were food deprived to 85% body weight and maintained at this weight throughout the experiment. Implanted animals were trained on a 2.64m Med Associates linear track with pellet dispensers at either end. Animals learned to run back and forth between the sides of the track and received a grain pellet at alternate ends. Each pellet received was considered a trial. The fasted animals were able to achieve over 120 trials in 30 minutes. (Animal were fed after daily training to maintain weight.) Six sessions were selected for analysis for each animal, with first and last recorded sessions automatically included. When not being run, animals were housed in individual cages with a 12hr light-12hr dark light cycle.

Six sessions from each animal were selection for analysis. Although each animal was run for at least seven sessions, occasionally an animal would refuse to run or run slowly (often the day following a run with a lot of pellets received). For the animal run the least number of times, only six sessions exceeded 50 trials per run. For consistency across analysis, we therefore selected the six sessions with the most trials to analyze from each animal. On average, the four animals ran 100.3+-33.1 trials per session, with the individual animals averaging 94.7+-35.3, 85.5+24.0, 114.5+-19.8, and 106.2+-41.2 trials per session.

Because the runs with the most trials were selected for analysis, trials were not always consecutive. If the first analyzed session is considered day 1, the analyzed sessions were as follows for each animal (as numbered by days on a calendar; animals were not necessarily run every day due to scheduling and equipment troubleshooting):

*Animal 1*: Days 1, 2, 3, 7, 8, 9 (*run across 9 days*)
*Animal 2:* Days 1, 2, 6, 7, 9, 13 *(run across 13 days)*
*Animal 3:* Days: 1, 2, 11, 14, 15, 16 *(run across 16 days)*
*Animal 4:* Days: 1, 2, 10, 14, 15, 16 *(run across 16 days)*

#### Calcium Imaging

One photon calcium imaging was done using UCLA V4 Miniscopes (Silva, 2017; Aharoni and Hoogland, 2019a), assembled with standard lens configurations (two 3mm diameter, 6mm FL achromat lens used in the objective module and one 4 mm diameter, 10mm FL achromat lens used in the emission module). We tested Ca2+ recording when one of the objective module lenses was replaced by a 3mm diameter, 9mm FL achromat lens, but no improvement in cell visualization was noted. Gain was set at high and Miniscope LED intensity varied between 10 and 38.

> Notes differentiating this method from mouse Ca2+ imaging: Base fluorescence appears lower in rats as compared to mice. While we have been advised that, using Miniscopes, gain can often be on “low” for mouse imaging, we found it necessary to have gain on “high” to visualize the highest number of cells.

#### GCaMP Visualization in Slice and Processing of Tissue Sections

Following conclusion of the study, animals were anesthetized and perfused with PBS then 10% formalin. The brain was then sliced into 60 micron coronal sections. For visualization as seen in Figure 1c, sections were mounted, let dry, and coverslipped with Vectashield. Sections were then imaged with a confocal microscope.

For GCaMP and PV+ visualization in Figure 1d, all solutions were prepared in phosphate buffered saline (PBS) containing 0.05% sodium azide and 2% normal donkey serum. Free-floating sections were rinsed in PBS three times for a total of at least 30 min between incubation steps. Tissue sections were sequentially placed in (1) 0.2% Triton X-100, 90 min; (2) mouse monoclonal anti-parvalbumin 2000X (Millipore), 48 hours; (3) rabbit anti-GFP 2000X,(AbCam), 24 hours; (4) biotinylated goat anti-rabbit serum 200X (Jackson Immunochemicals), 90 min; (5) a cocktail containing Cy3 conjugated donkey anti-mouse serum 200x (Jackson Immunochemicals) and FITC conjugated avidin 200x (Jackson), 90 min. Sections were coverslipped with a 9:1 mixture of glycerine and PBS containing 0.2% p-phenylenediamine and examined with a confocal microscope.

For cresyl violet visualization in Figure 1d, we used 0.1% cresyl violet solution containing 1% acetic acid. Sections were sequentially placed in (1) Histoclear, 5min; (2) 100% alcohol, 2min; (3), 95% alcohol, 1min; (4) 70% alcohol, 1min; (5) distilled water, 2min; (6) cresyl violet solution, 12min; (7) distilled water, quick rinses; (8) acid alcohol; 1.5min; (9) distilled water; 0.5min; (10) 70% alcohol, 0.5min; (11) 95% alcohol, 0.5min; (12) 100% alcohol, 0.5min, (13) Histoclear, 2min. Sections were then immediately coverslipped with Permount and examined with a confocal microscope.

### QUANTIFICATION AND STATISTICAL ANALYSIS

Means are expressed as mean ± the standard deviation. Overall averages are computed by finding average per animal then averaging those values. All analysis code is available at https://github.com/hsw28/ca_imaging.

#### Position and speed analysis

Position was sampled by an overhead camera at 30Hz. Position tracking was done post-recording using Bonsai (Lopes et al., 2015). Speed was determined by taking the hypotenuse of the coordinates of the point one before and one after the time point of interest. Speed was then smoothed using a Gaussian kernel of 1s and speed was converted from pixels/s to cm/s.

#### Video pre-processing

Videos were recorded with Miniscope software at 15 frames/second and 752 by 480 pixels. All video processing was done using the Matlab-based CIATAH software (Ahanonu, 2018; Corder et al., 2019) and processing files are available at https://github.com/hsw28/ca_imaging/. Videos were down sampled in space (3x using bi-linear interpolation) and time (2x). The video was normalized by subtracting the mean value of each frame from the frame. Each frame was then spatially filtered (normalized) using a bandpass FFT filter (70-100cycles/pixel). The video was then motion corrected to a chosen reference frame using TurboReg (Thevenaz et al., 1998). Videos were then converted to relative florescence (dF/F_0_), where F_0_ was the mean over the entire video.

#### Cell identification

Cells were first automatically identified using the Matlab-based CIATAH software (Ahanonu, 2018; Corder et al., 2019), using the CNMF-E method (Zhou et al., 2018). Processing files are available at https://github.com/hsw28/ca_imaging/. Images were filtered with a gaussian kernel of width 3 pixels. Neuron diameter was set at a pixel size of 13. The threshold for merging neurons was set at a calcium trace correlation of 0.65, and neurons were merged if their distances were smaller than 2 pixels and they had highly correlated spatial shapes (correlation>0.8) and small temporal correlations (correlation <0.4).

All cells identified using CNMF-E were then scored as neurons or not by a human scorer. Scoring was also done within the Matlab-based CIATAH software (Ahanonu, 2018; Corder et al., 2019) in a Matlab GUI. Human scoring was done while visualizing an activity trace, average Ca2+ waveform, a cropped movie montage of the candidate cell’s Ca2+ events, and a maximum projection of all cells on which the candidate cell was highlighted. The relative fluorescence (ΔF/F_0_) local maxima of each identified cell were considered calcium event times.

#### Place cell identification and spiking characteristics

Place cells were identified using two criteria. First, cells must have a mean firing rate greater than 0.01Hz during periods of movement (>12cm/s) (Wirtshafter and Wilson, 2019, 2020). Second, the mutual information (MI) computed in either direction of travel must be greater than 95% of MI scores computed 500 times from shuffled cells (Kinsky et al., 2018). To compute MI, the track was divided lengthwise into 4cm bins, the firing rate of each cell and the occupancy of the animal were found for each bin. Rate and occupancy were smoothed with a 2cm Gaussian kernel. Mutual information was computed during periods of movement in each direction of travel as follows (Olypher et al., 2003; Kinsky et al., 2018; Wirtshafter and Wilson, 2020):

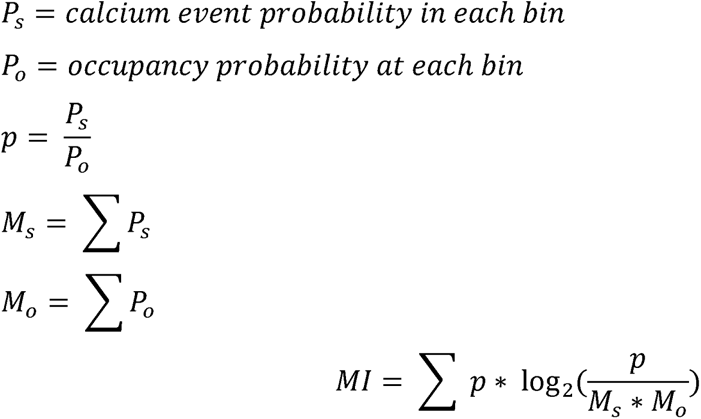

Place field location was determined by the location of maximum firing rate after binning position into 4cm bins. Speed modulation for individual cells was determined by computing average event rate when the animal was moving <=5cm/s and >=12cm/s; a cell was considered speed modulated if the rate at >=12cm/s was larger. At the population level, speed modulation was determined by examining population event rate versus speed. Cross correlations between speed and event rate were computed by binning speed and rate into 100ms bins.

#### Position decoding

Position was decoded using all recorded cells in a session and was decoded for times when the rat was moving (velocity>12cm/s). The track was split in 10cm segments and the time window of decoding was 0.5 seconds with non-overlapping time bins. Decoding was done using Bayesian decoding (Zhang et al., 1998).

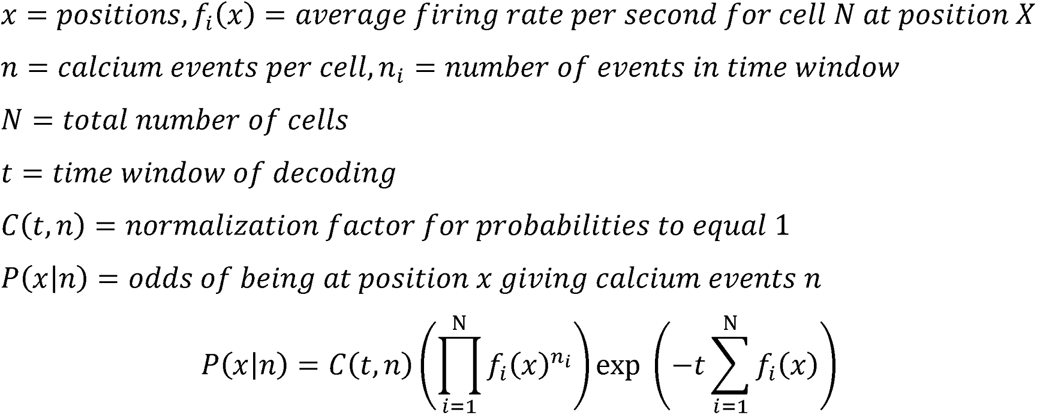

Accuracy decoding was determined by randomly shuffling firing per position bins and re-decoding. The difference between actual and decoded position was then determined per time bin and compared to decoded shuffled data.

#### Validation across days and within session

Validation was done using the Matlab-based CIATAH software (Ahanonu, 2018; Corder et al., 2019). Videos underwent 3 rounds of registration comprised of Turboreg image rotation (Thevenaz et al., 1998). An image binarization threshold of 40% of the images’ maximum value was used to remove background noise and axons and dendrites. A distance threshold of a maximum of 5 pixels was used to match cells across sessions, with a minimum 2-D correlation coefficient of 0.5. All sessions were aligned to the fourth recording session.

## Results

To initiate Ca2+ imaging, we injected AAV9-Syn-GCaMP7c below the CA1 cell layer in five male Fischer 344 x Brown Norway rats (Figure 1a, see Materials and Methods) (Nathanson et al., 2009). After injection, in the same surgery, cortex and corpus collosum were aspirated through a craniotomy until the alveus was clearly visible (Figure 1b, see Materials and Methods). A 2mm diameter GRIN lens was inserted into the craniotomy and cemented in place. Three animals were checked for GCaMP expression four weeks after surgery, and all three animals showed expression and were subsequently affixed with a base plate to secure the position of the Miniscope. The other two animals were checked six weeks after surgery and showed expression and were base plated. After the completion of the experiment, animals were sacrificed and expression of GCaMP in the CA1 pyramidal cell layer was verified (Figures 1c-d). Based on a comparison with cresyl violet staining of the hippocampal region, cellular GCaMP expression appeared to be located primarily in the CA1 principal cell layer (Figure 1d). Although contributions of interneurons to this study cannot be ruled out, antibody staining for parvalbumin positive (PV+) interneurons revealed that only a minority of GCaMP cells in the stratum pyramidale expressed parvalbumin (Figure 1d), consistent with observations that ∼3% of cells in the pyramidal layer are interneurons (Aika et al., 1994).

Animals were then food deprived to 85% body weight and trained on a 2.64m linear track (Figure 1e). Animals were rewarded with grain pellets on alternating ends of the linear track and quickly learned to run back and forth on the track. A visit to one reward well signified a trial. Animals were run for a minimum of 30 minutes or 50 trials, whichever came later, with total track time not to exceed one hour per session. Cells were recorded for 7 days for one animal, 11 days for one animal, and 16 days for two animals (n=4 rats, one of the five animals was eliminated when the implant was lost).

The six sessions with the most trials run were analyzed from each animal (average trials run = 100.3±33.1, see Materials and Methods), with first and last recorded sessions automatically included; sessions were not always consecutive (see Materials and Methods). These sessions were analyzed for cellular activity using the CNMF-E method implemented by the CIATAH software package (Ahanonu, 2018; Zhou et al., 2018; Corder et al., 2019) (See Materials and Methods). A total of 5761 neurons were recorded over the 24 sessions, with a maximum of 428 cells from a single session (mean of 239.6±90.0 cells) (Figure 2a-d). The number of cells recorded from each animal across sessions was relatively constant (Figure 2c).

**Figure 2:**
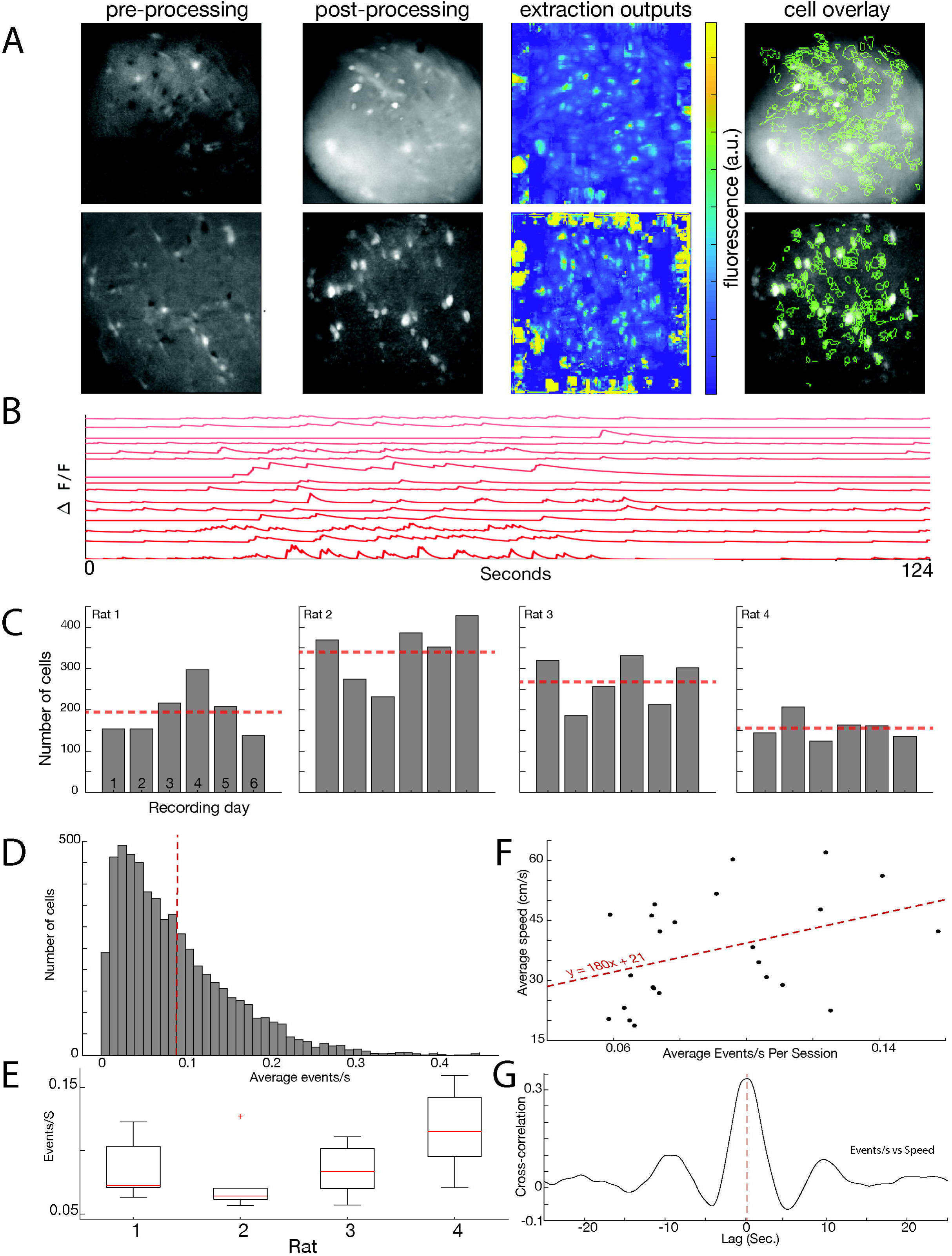
Hundreds of cells can be captured in a single Ca2+ imaging session. a. Examples of data from two (top and bottom rows) recording sessions. First column: example image from pre-processing. Second column: image post-processing (see materials and methods). Third column: all cell extractions, prior to manual cell sorting. Fourth column: post processing image overlayed with cell outlines identified after manual sorting. b. Example traces from 15 cells taken during 124 seconds of calcium imaging. c. Number of cells imaged per session for each rat. Dotted red line indicates the average for each animal. d. Histogram showing average rate (events/second) for individual cells. Dotted red line indicates the average (0.090+-0.019 events/s) e. The average calcium event rate (events/second) for individual rats (averages: 0.084+-0.024, 0.074+-0.026, 0.085+-0.020, and 0.116+-0.032 events/second) f. Correlation of average calcium event rate with average animal running speed. Average events per second in a session is positively correlated with the animal’s average speed in that session (p<0.05). g. An example from one session for the cross-correlation at different lags between the number of calcium events that occur in a second and the animal’s speed. In this example, the maximum cross correlation value at 0s was about 0.33, and the maximum value occurred at about 0.1s. On average, the cross-correlation at a lag of 0s was 1.8+-0.08, although the average maximum cross-correlation was 0.2 +-0.7 at an average lag of +0.47+-0.54s, indicating that calcium events reliably followed changes in speed, as opposed to vice versa.

We then determined average calcium events/s across animals to be 0.090+-0.019 events/s, with individual animals averaging 0.084+-0.024, 0.074+-0.026, 0.085+-0.020, and 0.116+-0.032 events/second. (Figure 2D-E). Some difference in average calcium event rate could be accounted for by differences in the base line speeds of the animals: there was a significant positive correlation between the animal’s average speed and the average events/s for each session (F(22)=4.2, p=0.05) (Figure 2F). Because speed is highly variable on a moment-to-moment basis, we also completed a cross-correlation between event rate and animal speed. In every session, we saw a positive cross-correlation between the animal’s speed and the number of calcium events across the population. The average cross-correlation at a lag of 0s was 1.8+-0.08, although the average maximum cross-correlation was 0.2 +-0.7 at an average lag of +0.47+-0.54s, indicating that calcium events reliably followed changes in speed, as opposed to vice versa (Figure 2G).

Significantly, calcium activity appeared to be heavily modulated by the animal’s activity, with periods of higher activity when the animal was running rather than stationary on the track (Figure 3). The biggest contributor to a change in event rate was transition from no movement to movement, with very few events occurring during periods of no movement (Figure 3C).

**Figure 3:**
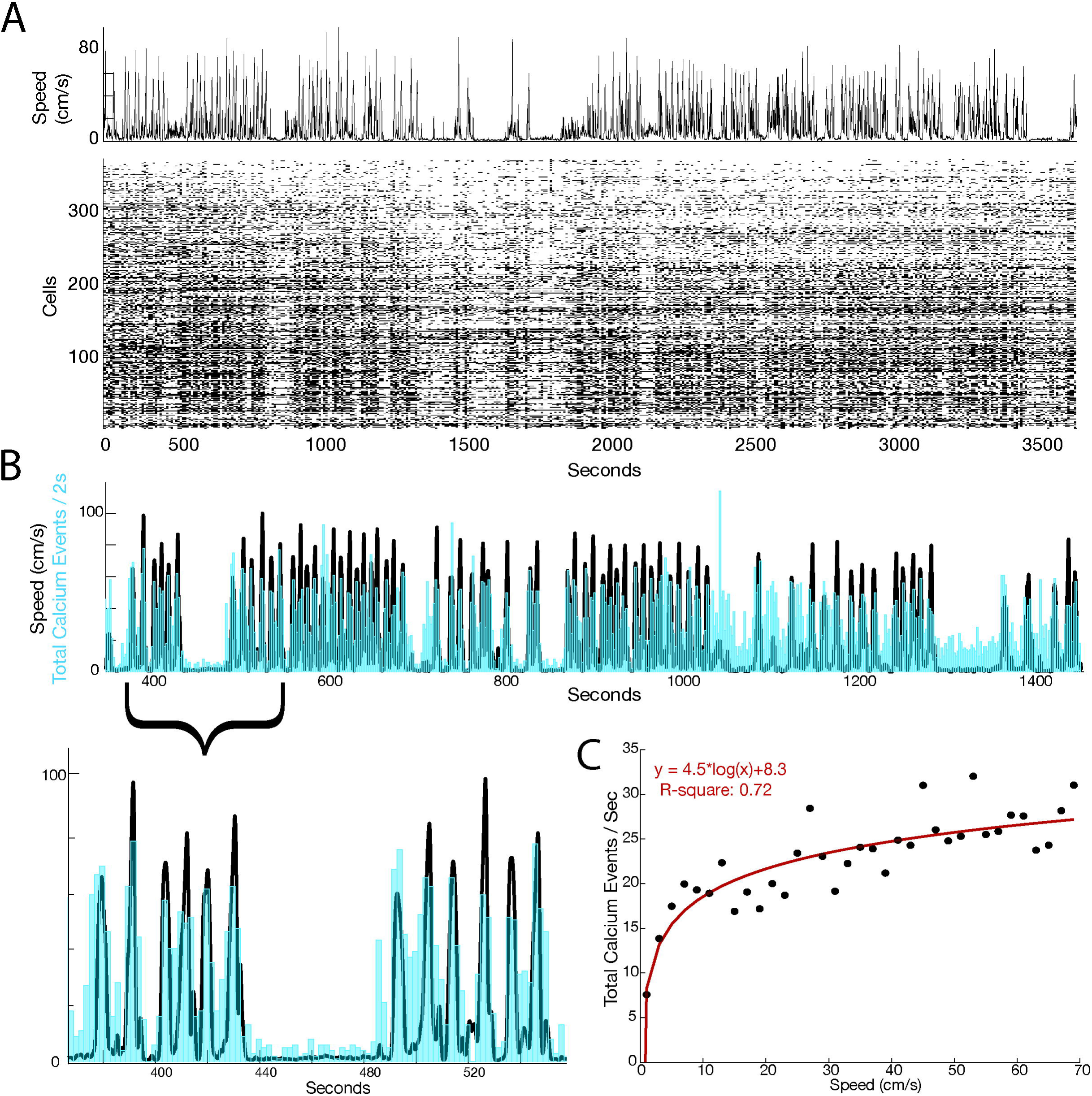
Calcium events are correlated with periods of movement. a. Raster plot of cell firing during linear track running. Top: the animal’s speed. Bottom: raster of peak calcium transients of 352 identified cells. b. Graph showing an example of speed (plotted in black) overlayed with event rate (in blue). Bottom panel is a zoom in on a smaller time period. c. An example logarithmic correlation of animal speed against calcium events/ second.

We then determined if there were place cells on any of the trials. A cell was considered to be a place cell if the mutual information (MI) computed in either direction of travel was greater than 95% of MI scores computed 500 times after shuffling the times of the calcium events that occur when the animal is moving (Ziv et al., 2013; Kinsky et al., 2018). Based on this criterion all trials contained place cells (Figure 3a). An average of 77.3±5.0%, of all cells recorded on the track were place cells (Figure 3), with individual rat averages at 71.6±10.6%, 77.3±1.2%, 76.4±9.5%, and 83.7±7.1%. We additionally determined the percentage of place cells if we only analyzed cells that had an average calcium events rate of greater than 0.01hz (Stefanini et al., 2020). This additional criterion yielded an even higher percentage of place cells, with 88.3±4.4% of these cells being classified as place cells. (Figure 3a).

We extended these results using an additional criterion previously used to determine the percentage of place cells using calcium imaging (Stefanini et al., 2020). We determined the percentage of cells that would be place cells if the mutual information value exceeded the mean plus three standard deviations of a shuffled distribution. Using this criterion, we found that 64.3+-5.1 percent of all cells were place cells, with individual animals at 59.5+-12.0%, 65.5+-2.3%, 61.2+-14%, and 71.0+-12.1%. If we applied the 0.01hz threshold for cell inclusion to this group, 73.4+-4.3% of cells were place cells, with individual animals at 68.0+-11.8%, 77.6+-5.6%, 77.0+-14.6% and 76.0+-13.0%.

We wished to further investigate the properties of these place cells and to confirm that cells were increasing their firing rate in place fields as is typical for pyramidal cells; as opposed to interneurons that mark spatial location by a substantial decrease in firing (Ego-Stengel and Wilson, 2007; Wilent and Nitz, 2007; Dombeck et al., 2010)). We examined the place cells’ maximum event rate during movement (presumably occurring in the place field) with their average firing rate during movement. The mean maximum rate was 0.15events/s, and the mean average rate was 0.02events/s. On average, cells increased firing 7.4x from their average firing rate during movement to their maximum firing rate (Figure 4D). We next determined if speed modulation occurred in the cells which display place fields. We compared whether speed-modulated neurons increased their in-field firing rate as much as non-speed-modulated neurons. We found no significant difference in firing rate increases between speed-modulated and non-modulated neurons (t-test (4023)=0.7102, p>0.05), nor were the distributions of increases different (KS-test p>0.05): both speed-modulated and non-modulated cells increased their firing rates, on average, >7x from baseline at the center of their place fields.

**Figure 4:**
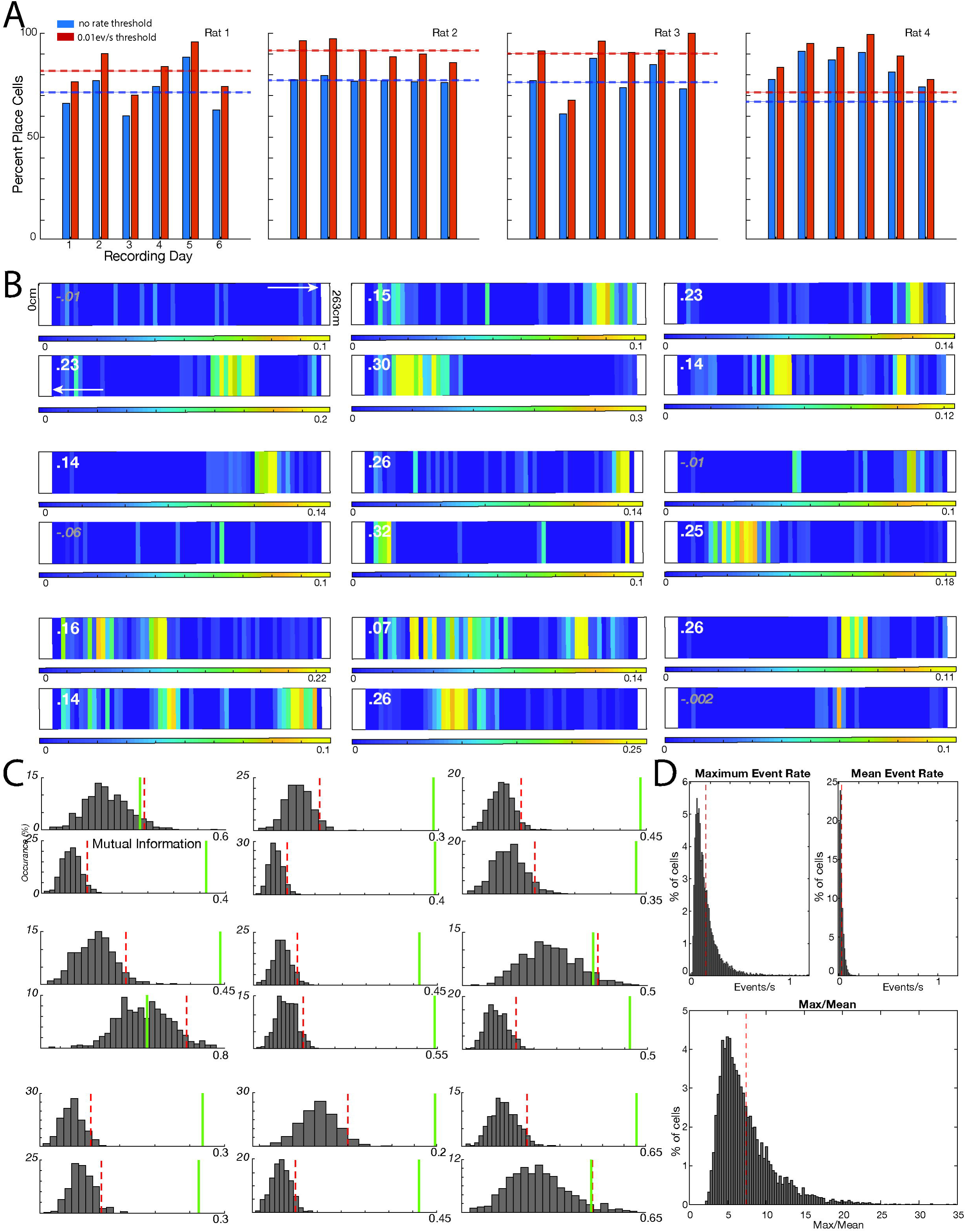
A high percentage of place cells are recorded on the linear track using Ca2+ imaging. a. Graph showing the percentage of place cells per session for each rat (blue). Analysis was repeated including only cells with a firing rate >0.01Hz during movement (red). Dotted lines indicate the average for each animal. b. Nine example place cells. A firing rate map in both directions of travel is provided for each cell, with the top map corresponding with running to the right, and the bottom map corresponding with running to the left. Note the colorbar scales may be different for each direction of travel. The difference in mutual information scores between the cell and the top 95% of shuffled data (actual MI-shuffled MI) is indicated in the left corner. If the actual MI was greater than 95% of shuffled data, the number is printed in bold white. c. Distribution of directional mutual information scores after shuffling cells seen in figure 3b. Event data was shuffled 500 times and the top 95% of mutual information scores were determined from shuffled data (red dotted line). If the actual mutual information score (green line) was greater than the upper 95% MI score of shuffled data, the cell was considered to be a place cell. d. Top left: Place field maximum event rate during movement (dotted red line indicates average of 0.15events/s). Top right: average firing rate during movement (red line is average at 0.02events/s). Bottom: Increase from average to maximum rate (max/mean). On average, cells increased firing 7.4x from their average firing rate during movement to their maximum firing rate (dotted red line indicates average)

We then aimed to determine whether place cell firing was sufficient to decode the position of the animal. We split the track in 10cm segments and used Bayesian decoding to determine the position of the animal in non-overlapping 0.5 second increments (Zhang et al., 1998). We found that the best positional decoding was accurate to a median error of 4.3cm (1.6% of the 2.64m track). Across all animals, accuracy ranged from a median of 4.3cm to 13.6cm (decoding errors were below 10cm in 23 sessions and above 10cm in one session) with a median error of 5.9cm (Figure 5a). To confirm that decoding accuracy was better than chance we shuffled firing rate per position bins and, using the shuffled data, decoded position. Median error for shuffled data ranged from 59.1cm to 77.6cm, which was significantly worse than non-shuffled data (Kruskal-Wallis test, x^2^(47)= 35.3, p=2.9*10^-9^) (Figure 5b).

**Figure 5:**
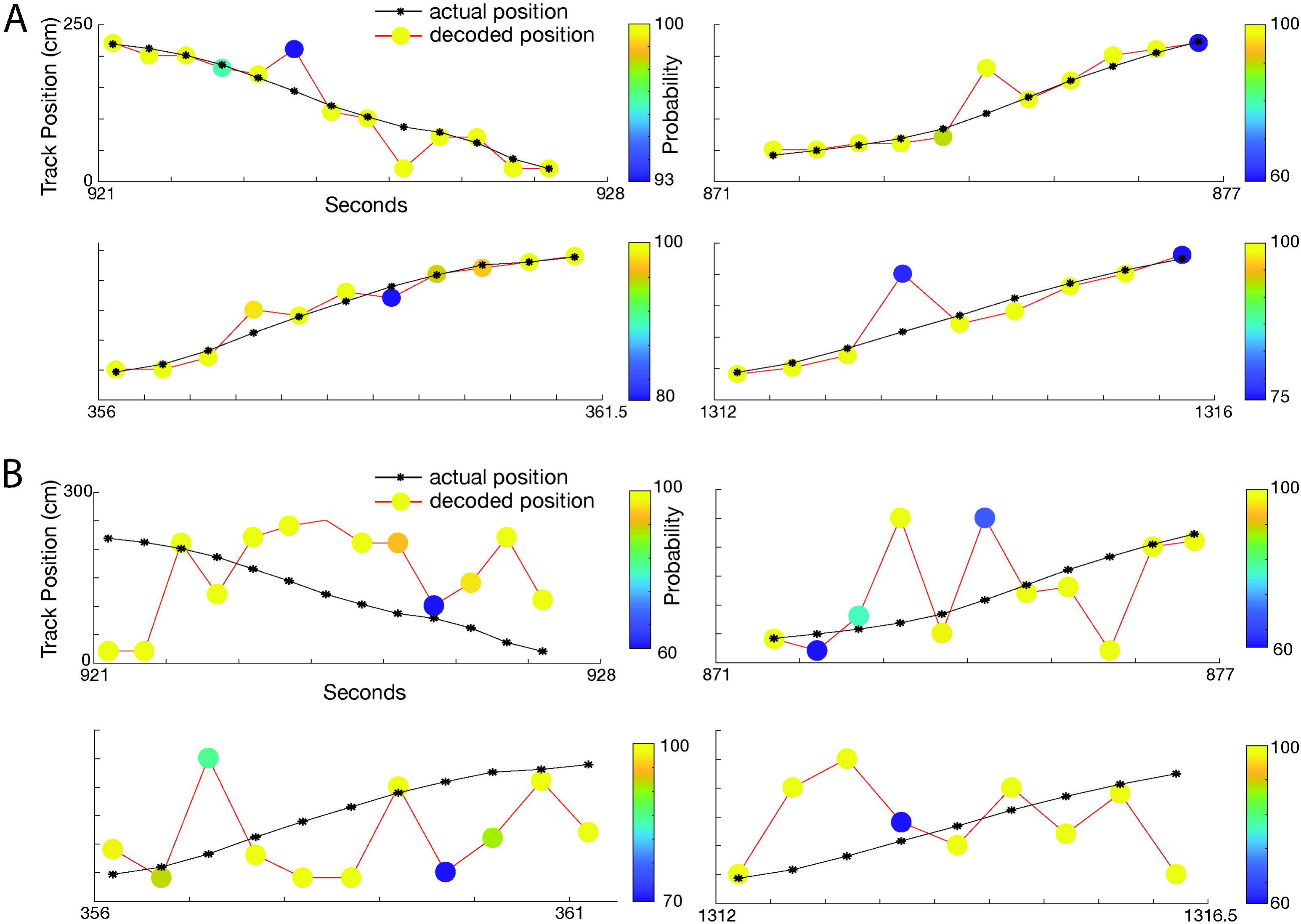
Place cells are sufficient to decode the animal’s position. a. Four decoding examples as the animal traverses the track. Black line indicates the animal’s actual position, with the stars indicated sample point. The colored circles connected by the red line indicate decoded position. The color of the circles indicates decoding certainty. b. Same as A but decoding is using shuffled units. Decoding is much less accurate using shuffled data.

We then aimed to determine if the same cells could be held over the duration of our recordings (six recording sessions, spanning 7, 11, or 16 days). 40% of the cells imaged on day one of the 7-day session were also present on day seven. Of the rat run for 11 days, 31% of the cells present on day one were also present on day eleven. For the two rats run 16 days, 28% and 50% of cells, respectively, that were present on day one were also present on day sixteen. Across all four animals, cells that were imaged on day one of recording had, on average, a 72±6% chance of being imaged on at least one subsequent day of recording (Figure 6A). (Of note, while the camera was placed in the same approximate position each day, thus allowing landmarks (such as blood vessels) to be followed throughout the study, we did not attempt to perfectly align landmarks or focal planes across days; instead, recordings each day were made to optimize total cell number in each day as proof of method.)

**Figure 6:**
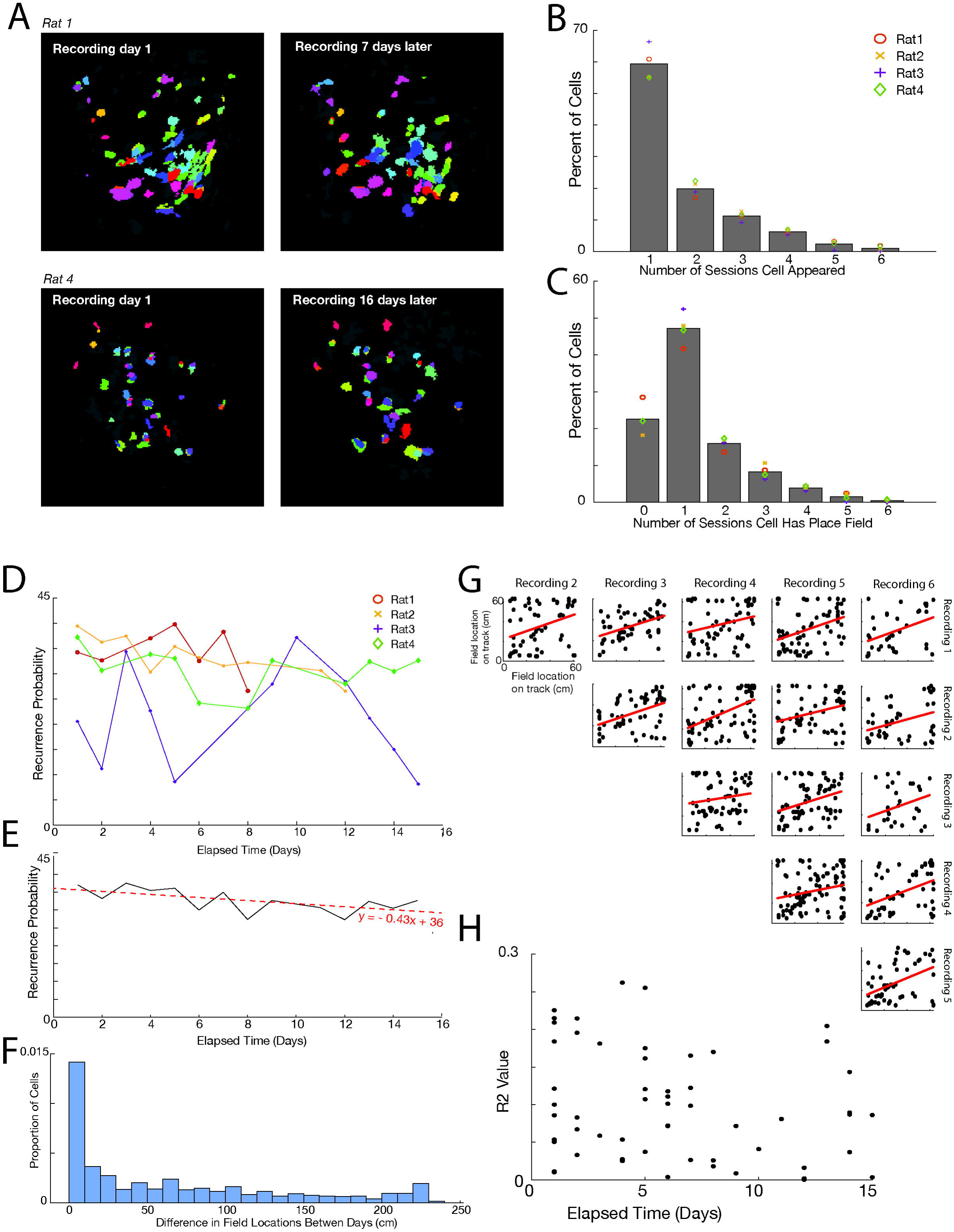
Individual cells and their place fields can be maintained across sessions. **a**. Cells can be maintained across sessions weeks apart. Examples from two animals of data taken on the first and final days of recording. Cells are color-matched from the first to last day. Cells in gray represent cells with no matches between the two days. Of note, while the camera was placed in the same approximate position each day, thus allowing landmarks (such as blood vessels) to be followed throughout the study, we did not attempt to perfectly align landmarks or focal planes across days; instead recordings each day were made to optimize total cell number in each day as proof of method. **b**. Percent of cells that appeared in different numbers of sessions. Symbols represent averages for individual animals. There was no significant difference in distribution of values between animals (non-parametric k-sample Anderson-Darling test, rank statistic: -1.48, p>0.05). c. Percent of cells that had place fields in different numbers of sessions. Symbols represent averages for individual animals. There was no significant difference in distribution of values between animals (non-parametric k-sample Anderson-Darling test, rank statistic: -1.59, p>0.05). **d**. The recurrence probability of a cell based on the number of days between recordings. Recordings from rat 3 were from found to be from two different focal planes. **e**. Average recurrence probability across days, excluding rat 3. For every day passing, a cell has a 0.43% less chance of reoccurring. **f**. Change in place field location between recording sessions. Included all sessions recorded for an animal; sessions did not need to be adjacent. g. A comparison of place field locations for one rat across all six recording sessions (15 comparisons). For all 15 comparisons using this animal, place field locations were highly correlated between the sessions. Each point indicates the place field of a single cell identified in the two sessions indicated. Red lines on the graphs indicate a p<0.05 using an F-test. Across all four animals, we found field locations to be consistently correlated, with 93% of session comparisons having a significantly positive correlation between place field locations (f-test, p<0.05). **h**. There was no correlation (p>0.05) between r2 value of place field correlations and days elapsed between sessions.

We then looked to determine how often a cell appeared in each session. Across all cells, 59.4+-5.4% of cells were visible in only one session, 19.9+-2.4 were visible in two, 11.2+-1.5 in three, 6.3+-0.8 in four, 2.3+-1.3 in five, and 1.0+-0.9 were in all six sessions (Figure 6B). We then looked at place cells that were repeated across sessions. As stated, 22.6+-4.3% of cells did not have place cells in any session. Of the 77.4% of cells that did have place fields, 61.0+-5.7% (47.2% of total cells) had fields in only one session, 20.6+-2.0 (15.9% of total) had fields in two sessions, 10.8+-2.4% (8.3% of total) had fields in three sessions, 5.1+-0.7% (4.9% of total) had fields in four sessions, 2.0+-1.4% (1.5% of total) had fields in five sessions, and 0.5+-0.5 (0.4%of total) had fields in all six sessions (Figure 6C). Place cell recurrence was not statistically different from the recurrence of all cells (paired t-test(5)=2.04*10^-15^, p=1).

We then determined the recurrence probability of a cell based on the number of days apart the recording took place (Figure 6D). Three of our animals had relatively consistent recurrence probabilities, while one animal (animal 3) was highly variable. Going back to the recordings, we saw that recordings from this animal came from two different focal planes, causing the recording of two different cell populations. Excluding this animal, there was a modest trend across days for a decreasing recurrence probability, with the probability of a cell reoccurring decreasing by 0.43% per day separating the recording periods (Figure 6E).

We then determined whether a cell’s place field remained in the same location during days when the place cell reoccurred. We found that, across all days of recording, the mean standard deviation of place field locations was about 36cm (∼13% of track length), with the individual animals at 39cm, 36cm, 32cm, and 37cm. We compared these distances with distances if place field locations were shuffled, and we repeated shuffling 500 times. All four animals had a smaller deviation between fields than would be expected by chance (p<0.05 for all animals). Between individual days (including nonadjacent days), the change in field distance was highly variable at 65cm+-71cm, with a median of 40cm and a large concentration of values around 0cm (Figure 6F).

Although fields across days were more consistent than would have been expected by random chance, we had not yet determined whether the animal had a consistent map across all recording days (Ziv et al., 2013), or if it switched between multiple maps (Sheintuch et al., 2020). To determine this, we compared place field location for cells in a session against the place field locations of the same cells in another session. To do this, for each cell, we plotted the location of the place field for that session against the place field location of the same cell for another session. We did this comparison for every cell that had a place field in more than one session. If place field mapping was similar from session to session, we would expect the plotted field locations of one session to have a strong positive correlation with the locations from another session. Of the pairs of sessions that had more than 10 place cells shared between sessions (57 out of 60 total session pairs—15 session pairs for rat 1, 15 for rat 2, 14 for rat 3, and 13 for rat 4), 53 of these pairs (93%) had a significant positive correlation (f-test, p<0.05) between the place field locations, with no evidence of switching between maps across days (Figure 6G). We found that correlations between field locations did not decay as a function of the number of days between sessions (Figure 6H).

## Discussion

In this manuscript, we have demonstrated calcium imaging recordings can be performed in the CA1 region of the hippocampus in freely moving rats. These recordings can capture hundreds of cells in a single recording session, including a high number and percentage of place cells sufficient to decode the animal’s position to within centimeters of accuracy, and that place cell field location is consistent across days. Calcium imaging in rat will allow for more behaviorally complex and translational experiments than have been done in mouse.

### Measuring Spatial Content and Categorizing Place Cells

We chose to use mutual information as the measure of spatial content for multiple reasons: First, it has precedence for use in both calcium imaging (Ziv et al., 2013; Meshulam et al., 2017; Kinsky et al., 2018; Kinsky et al., 2020; Stefanini et al., 2020) and electrophysiology studies (Jun et al., 2020) (Table 1). Second, threshold measures commonly used in electrophysiology that depend on firing rate (such as “a cell is a place cell if firing rate, as measured by electrophysiology, exceeds 2Hz for a 10cm area”) (Rich et al., 2014) (Table 1) do not transfer well to calcium imaging experiments as, using currently available GCaMPs, calcium events are captured much less frequently than spikes in electrophysiology (typically, ∼0.02-0.06 events/s is the mean rate observed during 1P calcium imaging, versus >0.5-1.5 spikes/s in electrophysiology, a 25-fold difference, see Table 1). Third, other measures of information content, such as bits/spike or bits/second (Table 1), tend to be highly sensitive to changes in firing rate, including baseline firing rate (Souza et al., 2018), as compared to mutual information. Specifically, cells with lower baseline firing rates are favored to have higher baseline bits/spike or bits/second values (Souza et al., 2018; Wirtshafter and Wilson, 2020), which makes it not a useful metric with which to compare values from calcium imaging with data collected from electrophysiology. Additionally, mutual information is advantageous in that it is a better predictor of decoding accuracy than bits/spike and bits/second, and thus may better capture spatial informational content (Souza et al., 2018). The criteria we used to identify place cells were as or more stringent than previously used criteria in calcium imaging that identified a significantly lower percentage of place fields (Table 1).

**Table 1.** Comparison of place cell findings in previous studies in mice and rats

| Study | Method | Species | Task | Mean spike rate or Ca event rate | % place cells | How place cells were determined |
| --- | --- | --- | --- | --- | --- | --- |
| Guzowski et al. 1999 | IEG | Rat | Cylinder and platform exploration | NA | **38-39%** | ARC detection |
| Witharana et al. 2016 | IEG | Rat | Environmental exposure and circular track | NA | **22-73%**, mean 33±18% | Homer1a and ARC detection |
| Talbot et al. 2018 | Ephys | Mouse | Open field | 1.14±0.12 spikes/s for all pyramidal cells<br>1.49±0.15 spikes/s for place cells<br>2.61 ± 0.2 spikes/s mean in-field rate | **47.8%** | overall discharge rate was 0.1-2 AP/s, the spatial coherence was >0.4, and the information content was >0.4 bits/spike |
| Jun et al. 2020 | Ephys | Mouse | Open field and linear track | ~0.8 spikes/s mean rate<br>~3.5 spikes/s maximum rate | **50%** | Mutual information >95% of shuffled data |
| Maurer et al. 2006 | Ephys | Rat | Circular track | Not noted | **~43%** on smaller track<br>**~68%** on larger track | place field has a single cycle (360%) phase precession |
| Wirtshafter and Wilson 2020 | Ephys | Rat | Double sided T maze | ~0-5hz in field | **75.2%** exceeded 0.8 bits/spike<br>**63%** had at least one place field and exceeded bits/spike threshold | 1) Bits/second >0.8<br>2) firing in a region must be greater than two standard deviations greater than the mean firing rate and place field area must be at least 15 cm long |
| Fenton et al. 2008 | Ephys | Rat | Forage for food in cylinder and | 0.91-1.01 spikes/s depending on | **64%** in smaller cylinder | cell coherence was >0.25 and information |
|  |  |  | chamber | environment<br>2-2.5 spikes/s<br>mean in-field<br>rate | **72%** in 6x<br>larger<br>chamber | content was >0.5<br>bits/AP, and if the<br>cell had at least<br>one place field. |
| Markus et al.<br>1994 | Ephys | Rat | 8 arm radial<br>maze forced<br>choice | 0.56 spikes/s<br>in young<br>animals | **37%** | Bits/spike cutoff of<br>2.29 |
| Rich et al.<br>2014 | Ephys | Rat | 48m long<br>track | Up to ~0.7<br>spikes/s<br>during<br>running,<br>highly<br>correlated<br>with # of place<br>fields | **65%** at 48m<br>(range 46-77%) | A place field was<br>defined as $\geq 15$<br>contiguous cm of<br>rate map in where<br>firing rate > 2 Hz. |
| Meshulam et<br>al. 2017 | 2p<br>calcium<br>imaging | Mouse | VR linear<br>track | Not noted | **~30%**, range<br>28% to 41%. | Mutual information<br>above 80% of<br>shuffled |
| Ziv et al. 2013 | 1p<br>calcium<br>imaging | Mouse | Linear track | ~0-0.5<br>events/s | **20%** | Mutual information<br>above 95% of<br>shuffled |
| Stefanini et al.<br>2020 | 1p<br>calcium<br>imaging | Mouse | Open field | 0.06 ± 0.04<br>ev/s | MI mean+3σ:<br>**~4%** for all<br>cells<br>**~10%** with<br>0.1ev/s cutoff<br>>95% shuffled<br>MI<br>**~12%** for all<br>cells<br>**~22%** with<br>0.1ev/s cutoff | Mutual information<br>> mean+3σ of the<br>shuffled<br>distribution.<br>Also compared to<br>mutual information<br>above 95% of<br>shuffled |
| Kinsky et al.<br>2018 | 1p<br>calcium<br>imaging | Mouse | Square and<br>octagon<br>tracks | Not noted | Unspecified | 1) had $\geq 5$<br>calcium transients<br>during the session<br>2) Mutual<br>information above<br>95% of shuffled |
| Kinsky et al.<br>2020 | 1p<br>calcium<br>imaging | Mouse | Continuous<br>alternation | 0.02-0.03 ev/s<br>(mean ~40-50<br>events in<br>~30m<br>session) | Unspecified | 1) had $\geq 5$<br>calcium transients<br>during the session<br>2) Mutual<br>information above<br>95% of shuffled |
| This work | 1p calcium imaging | Rat | Linear track | 0.02events/s during movement only<br>0.15events/s mean maximum in-field rate | >95% shuffled MI<br>**77.3%** for all cells<br>**88.3%** with 0.01ev/s cutoff<br>MI mean+3σ:<br>**64.3%** for all cells<br>**73.4%** with 0.01ev/s cutoff | Mutual information above 95% of shuffled<br>Also computed mutual information > mean+3σ of the shuffled distribution. |

Based on anatomical data (Figure 1d), it is reasonable to conclude that the cells we recorded from were primarily pyramidal cells in the stratum pyramidale. GCaMP expression appears to be most densely expressed in pyramidal cells in the cell layer (Figure 1d) (although dendritic processes can be viewed extending into the radiatum, where occasional cells are seen (Figure 1c-1d)). Further, many of the labeled cells displayed an obviously pyramidal morphology (Figure 1c) and only a small proportion of total GCaMP labeled cells expressed parvalbumin, a result consistent with reports that only about 3% of cells in the stratum pyramidale are interneurons (Aika et al., 1994).) The majority of double labeled cells were in the stratum radiatum, or on the border between the stratum pyramidale and the stratum radiatum, and would likely be too deep and too obscured to be captured by the implanted lens and Miniscope. (Even including the radiatum, double labeled cells constitute a minority of GCaMP labeled neurons).

In addition to anatomical and morphological data, the cells we imaged had calcium event patterns characteristic of pyramidal cells. It is well established that pyramidal place cells increase their firing rate in their place field, while interneurons that express place specificity do so by decreasing their firing rate. Both ways of specifying place presence would result in an elevated mutual information score. However, given that the vast majority of cells we observed displayed an increased rate of calcium events at specific locations (Figure 4d), it is likely that these cells are pyramidal excitatory cells. The percentage of place cells recorded in the present study are comparable to those seen in electrophysiology (Table 1), where interneurons can be more easily separated from pyramidal cells based on spike morphology and frequency, suggesting that we may be imaging from a similar population of cells.

These considerations suggest that any contribution of interneurons to the observed effects would, at most, be very small.

### Differences in place cell findings between electrophysiology, intermediate early gene studies, and calcium imaging

The percentage of hippocampal CA1 cells found to be place cells using calcium recordings (77.3+-5.0%) was also very slightly above the upper range of findings from electrophysiological data in rats, which report that 35-75% of CA1 pyramidal cells recorded on a navigational task are place cells (the number varies widely depending on task parameters, with studies using a linear track task reporting the highest number of place cells) (Table 1) (Markus et al., 1994; Fenton et al., 2008; Wirtshafter and Wilson, 2020). There are multiple possible reasons related to recording technology as to why our calcium imaging experiments detected a higher percentage of place cells than traditional electrophysiology experiments: the difference likely resulted from electrophysiology capturing (as compared to calcium imaging) both a lower number of cells with active place fields detected (a smaller numerator) and a larger number of cells without active place fields detected (a larger denominator). First, in electrophysiology experiments with moveable electrodes (such as tetrodes), electrodes are typically positioned while rats are inside a “sleep box” or other non-training apparatus until cells are detected. The animal is then placed on a track for experimentation. This practice optimizes cell detection for place cells that are active in the “sleep box” (both place cells with fields in the sleep box, and place cells that are reactivated during ripples during sleep), rather than cells that have place fields on the track. Adjusting and recording in a sleep box therefore increases the number of total cells recorded that do not have fields on the track and therefore reduces the place cell percentage. Because calcium imaging uses a fixed lens location and images a large number of cells at once, recording is not optimized for a particular environment. This may result in the recording of a higher percentage of active place cells on the recording apparatus. (Due to logistical constraints in this study, we did not complete any recording while the animal was not on the track.) Second, while an effort is made in electrophysiology to discard times on the track when the animal is not moving, and thus to remove corresponding spiking during ripples, this removal is never perfectly done. The recording of ripples during electrophysiology has two effects: the recording of cells that show replay on the track but do not have place fields on the track and, second, the recording of ‘extra-field’ ripple spiking from cells with fields on the track. This ‘extra-field’ spiking can cause place fields to seem less defined, and cause corresponding distortion of measures of spatial information, such as mutual information and bits/spike. This, in turn, may cause these cells not to be classified as place cells. Conversely, in calcium imaging, very few to no calcium events are picked up during periods of no movement (see later in discussion, figure 3), so both of these issues (increasing the count of cells without fields on the track, and the distortion of spatial information for place cells) are avoided. Third, while current GCaMPs, such as the GCaMP7c used in this study, have improved single spike resolution compared to past GCaMPs, they still fall short of the single spike resolution of electrophysiology (Tian et al., 2009a; Chen et al., 2013; Podor et al., 2015a; Podor et al., 2015b; Shemesh et al., 2020). This property of calcium indicators causes individual spikes, such as those that may occur on the track outside a cell’s place field, to be missed, therefore biasing calcium imaging towards fast cell activity that only occurs in the confines of a place field. Calcium imaging therefore, again, results in potentially higher spatial information metrics as compared to electrophysiology, which may cause the identification of more place cells. Finally, compared to electrophysiology, calcium imaging provides additional characteristics (cell shape, size, and brightness) by which to separate adjacent or nearby cells. Because conflation of multiple cells in electrophysiology may disguise place fields and disrupt place field measurements (such as mutual information or bits per spike), the additional resolution afforded by calcium imaging may allow for more accurate place cell differentiation and identification.

Similar to electrophysiology and in contrast to intermediate early gene (IEG) studies, calcium imaging is biased towards capturing cells active in an environment and not the total cellular population. Additionally, as discussed above, due to limits on single spike resolution, calcium imaging may be picking up even fewer cells with infrequent spiking than does electrophysiology. Thus, this bias of calcium imaging, and electrophysiology, toward detecting only active cells may yield a higher percentage of place cells than do intermediate early gene studies that estimate a total of 30-40% of cells to be active in any given environment (table 1) (Guzowski et al., 1999; Witharana et al., 2016).

### The percentage of place cells in rat calcium imaging exceeds the percentage in mouse calcium imaging

Significantly, we found that a higher percentage of cells were place cells in our calcium imaging experiments in rats than those reported in mouse CA1 calcium imaging studies (Table 1). This may, in part, reflect the fact that electrophysiological studies indicate that place cells are less frequent in mouse than rat (Table 1), a finding that may be related to reports that mouse place cells are less spatially tuned, contain less spatial information, and have less stable place fields than those in rat (Hok et al., 2016; Mou et al., 2018). More striking, however, is the fact that whereas the percentage of place cells observed in the current study is only slightly higher than that observed in rat electrophysiological studies, examination of Table 1 indicates that calcium imaging experiments in mice typically yield values drastically lower than those observed in mouse electrophysiological studies (mouse calcium imaging experiments have found place cell percentages as low as 4%) (Stefanini et al., 2020). These comparisons suggest that there is a much greater concordance between calcium imaging and electrophysiological studies in the rat than in the mouse. These discrepancies may reflect physiological differences between rats and mice that affect the type of activity captured in calcium imaging recordings. CA1 cells in mice are more excitable than those in rat (Routh et al., 2009; Hok et al., 2016), and prolonged GCaMP expression in mice has been shown to lead to cell hyperexcitability, seizures, and even cell death (Tian et al., 2009b; Grienberger and Konnerth, 2012b; Resendez et al., 2016; Steinmetz et al., 2017; Yang et al., 2018). The higher baseline excitability of mouse cells, combined with a further increase in cellular excitability after GCaMP expression may result in a lower threshold for cell firing and less defined place fields, less stable fields, and, therefore, a seemingly lower number of place cells compared to electrophysiology recordings (Hok et al., 2016). Because rats have a lower baseline excitability, they may be less prone to these GCaMP effects, although further slice electrophysiology work will need to be done to directly compare the effect of GCaMP on mouse and rat neurons.

We considered that differences in calcium event rate between our data and other studies might lead to the high percentage of place cells we saw in this work. On one end of the spectrum, Stefanini et al 2020 found a particularly low percentage of place cells (∼4% of recorded cells in mouse CA1, using calcium imaging) and investigated whether this might be a consequence of collecting many cells with low calcium event rates. Their reported mean rate was 0.06+-0.04 events/s as recorded in mice in an open field, consistent with 0.05-0.5 events/s in other calcium imaging experiments in mouse. Even after eliminating cells with <0.1events/sec (a 10x less stringent criteria than we used in Figure 3A), they still only found that 10-20% of cells were place cells (Stefanini et al., 2020). Conversely, Ziv 2013, who reported a high (for mouse) 20% of imaged CA1 mouse cells were place cells, reported a higher mean event rate, at ∼0.1-0.5 events/s (Ziv et al., 2013). Our results (0.090+-0.019 events/s, Figure 2D-E) are on the lower end of those reported in Ziv et al, and potentially consistent with the fact that rat neurons are somewhat less excitable than those in mice (Routh et al., 2009; Hok et al., 2016). Additionally, our very slightly lower event rate could be the result of less ‘out of field’ firing of place cells, which also tends to be more predominant in mice (Hok et al., 2016). However, our results are within the range of those reported in mouse imaging studies (Table 1), so differences in event rate are not a likely to be a major contributor to the place cell percentage we observed.

It is also possible that viral infection of different populations of cells may have contributed to the observed rat/mouse differences. For example, if mouse calcium imaging experiments were recording a larger proportion of interneurons than was the case in the present experiment, this might alter the proportion of place cells. However, in the numerous 1-photon calcium imaging experiments done of mouse hippocampus, we could find little documentation regarding the percentage of excitatory versus inhibitory neurons labeled (Ziv et al., 2013; Kinsky et al., 2018; Kinsky et al., 2020; Stefanini et al., 2020). Additionally, to our knowledge, little data on the location GCaMP expression in the mouse HPC are available, and, given the low total number of interneurons in the pyramidal cell layer, it seems unlikely that such an effect could be of sufficient magnitude to account for the observed species differences.

In addition to potentially allowing the recording of a higher number of place cells, the lower excitability of rat CA1 neurons, compared to mouse, also has other advantages. We saw no behavioral evidence of post-operative seizures in any of our implanted animals, and no behavioral or imaging evidence of seizures several months post viral injection and lens implementation. This is in contrast to calcium imaging experiments in mice, which frequently report that animals have to be removed from behavior or analysis due to the development of seizure activity (Steinmetz et al., 2017; Huang et al., 2021).

### 1P calcium imaging using GCAMP7c is movement responsive and minimally activated when the animal is still

Importantly, while calcium transients and events are correlated with action potentials, they are far from a 1:1 correspondence and may capture more or less information than electrophysiology (Huang et al., 2021). During our recording sessions, we saw clear evidence that calcium event rate was correlated with periods of movement (Figure 3). In electrophysiology experiments, place cell spiking is generally linearly correlated with running speed (McNaughton et al., 1983; Huxter et al., 2003), and significant spiking occurs during ripples when the animal is not moving (Ylinen et al., 1995; Chrobak and Buzsaki, 1996; Csicsvari et al., 1999; Kudrimoti et al., 1999). Our finding greatly differs from the patterns of spiking seen in electrophysiology data, as we saw minimal calcium events during periods of no movement, with a pattern of logarithmic growth of the number of calcium events as the animal began moving and sped up (Figure 3C). Because the vast majority of CA1 pyramidal cells are speed and/or ripple modulated, and we have imaged primarily pyramidal cells, this result is likely not explainable by the population of cells that express GCaMP (Huxter et al., 2003). Our result is consistent with calcium imaging experiments done in mice by Zhou et al., in which more calcium events were seen during movement in an environment where food reward could be obtained than during quiet wakefulness in a neutral box (Zhou et al., 2019). In our experiment, periods of quiet wake were in the same environment (the track) as periods of movement, which shows that changes in calcium event frequency are related to movement and not to contextual associations. Zhou et al. posit that the high amount of calcium activity that occurs during movement, but not during sharp wave ripples in mouse, implicate calcium events in theta dependent processes, such as memory encoding and planning (Buzsaki, 1989; Hasselmo et al., 2002; Hasselmo, 2005; Shirvalkar et al., 2010; Colgin, 2013; Wikenheiser and Redish, 2015; Zhou et al., 2019; Wirtshafter and Wilson, 2021). It is possible, and consistent with slice data, that NMDA receptors, which permit calcium influx, are minimally involved in cell spiking during ripples, and the process is more heavily modulated by AMPA receptors, which do not permit calcium influx (Maier et al., 2003; Colgin et al., 2004). There is also evidence that activity during ripples can be visualized using 2-photon calcium imaging in mice (Malvache et al., 2016; Grosmark et al., 2021), although we could not find any experiments showing this activity using 1-photon imaging. It is therefore unclear whether limits to imaging spiking activity during ripples is due to the imaging technique or to the use of rats. Further work will be needed to determine if, in rat, frequent calcium events are also unique to movement, as opposed to sharp wave ripples.

### Results show stable place cell identification and these cells have stable place fields over days

Our findings on cell activity over days were similar to and consistent with prior results reported in mice. Prior work has reported that about 57% of cells are active in only one or two sessions (Ziv et al., 2013); we found a higher proportion (59.4+-5.4% of cells were visible in only one session, 19.9+-2.4 were visible in two) (Figure 6B), likely because we worked to optimize cell count over precisely consistent focal planes. We also reported that probability of a cell reoccurring decreased by 0.43% per day separating recording sessions. This result was exactly consistent with prior results in mouse that cell reoccurrence decreases from ∼25% for sessions 5 days apart to ∼15% for sessions 30 days apart (25 days difference * 0.43 = ∼10.8% change)(Figure 6E) (Ziv et al., 2013). This potential cell turnover did not impact the accuracy of the representation of the environment, as decoding accuracy was consistent throughout sessions (Levy et al., 2021).

There are differing results regarding whether an environment has a single semi-stable map (Ziv et al., 2013; Mau et al., 2018; Kinsky et al., 2020), or there are multiple maps, involving the same cells, that code for a single environment (Sheintuch et al., 2020). In this study, we only saw evidence of the former, as the vast majority of sessions had cells with place fields correlated to previous sessions (Figure 6F) and place field locations did not change as a function of the number of days between sessions (Figure 6G). In other words, cell reoccurrence decreases across days, but apparently when a cell does reoccur, its place field remains largely the same.

## Conclusion

As previously mentioned, similarities between rat and human physiology, as well as the increased capacity for complex tasks of learning and memory (Whishaw, 1995; Frick et al., 2000; Whishaw et al., 2001; Cressant et al., 2007; Rosenfeld and Ferguson, 2014; Hok et al., 2016), position rats as a well-suited model organism for studying hippocampal function, with several advantages over the mouse. Rats are advantageous over mice in that, for many diseases and disorders, including those with substantial hippocampal involvement, their physiology and behavior more closely mirrors that of humans (Ellenbroek and Youn, 2016). For instance, 5-HT serotonin receptors, which are prominent in the hippocampus and are implicated in a variety of disorders (including mood and anxiety disorders, psychiatric disorders such as schizophrenia, addiction and ADHD), are distributed in rats, but not in mice, in a similar pattern as seen in humans (Hirst et al., 2003; Ellenbroek and Youn, 2016). Hippocampal neurogenesis, which is believed to play an important role in learning, memory, and the treatment of depression, is believed to follow a similar pattern in rats and humans, and is less pronounced in mice (Snyder et al., 2009). In addition, both pharmacologic and behavioral responses to addiction are more similar between rats and humans than mice and humans (Jupp et al., 2013; Parker et al., 2014; Ellenbroek and Youn, 2016).

Having the ability to do calcium imaging in rat CA1 will allow increased research into these disorders, as well as basic research in learning and memory. Because rats are substantially larger than mice, they can support larger lens implants and cameras, which may lead to the ability to record from thousands of hippocampal cells in a single session during complex tasks. The ability to record a large number of cells across days will allow research into cell firing changes in the hippocampus, which will provide insight into more advanced learning and disease mechanisms that are difficult to understand when only investigating mice. Additionally, the uniquely high percentage of recorded place cells (>80% of recorded cells) makes CA1 calcium imaging in rats an ideal method by which to study spatial navigation and learning over days, including mechanisms behind memory consolidation, spatial remapping, and planning over time.

## Acknowledgements

We would like to thank all members of the Disterhoft lab, especially Matthew Oh, Craig Weiss, and Kent Park. We would also like to thank the Miniscope team, in particular Daniel Aharoni and Federico Sangiuliano Jimka. We additionally thank Biafra Ahanonu for assistance with CIATAH, Amy Christensen for providing advice during imaging setup, Drew Ames for consultation on figures, and David Wirtshafter for discussion, editing, and his assistance and expertise with immunohistochemistry.

## Related Manuscripts

Authors do not have a related or duplicate manuscript under consideration (or accepted) for publication elsewhere.

## Citations

Ahanonu B (2018) CIAtah: a software package for analyzing one- and two-photon calcium imaging datasets. In, v1.0.0 Edition: Zenodo.

Aharoni D, Hoogland TM (2019a) Circuit investigations with open-source miniaturized microscopes: past, present and future. Frontiers in cellular neuroscience 13:141.

Aharoni D, Hoogland TM (2019b) Circuit Investigations With Open-Source Miniaturized Microscopes: Past, Present and Future. Frontiers in cellular neuroscience 13:141.

Aharoni D, Khakh BS, Silva AJ, Golshani P (2019) All the light that we can see: a new era in miniaturized microscopy. Nat Methods 16:11–13.

Aika Y, Ren JQ, Kosaka K, Kosaka T (1994) Quantitative analysis of GABA-like-immunoreactive and parvalbumin-containing neurons in the CA1 region of the rat hippocampus using a stereological method, the disector. Exp Brain Res 99:267–276.

Andrzejewski ME, Ryals C (2016) Dissociable hippocampal and amygdalar D1-like receptor contribution to discriminated Pavlovian conditioned approach learning. Behav Brain Res 299:111–121.

Anner P, Passecker J, Klausberger T, Dorffner G (2020) Ca2+ imaging of neurons in freely moving rats with automatic post hoc histological identification. Journal of neuroscience methods 341:108765.

Avigan PD, Cammack K, Shapiro ML (2020) Flexible spatial learning requires both the dorsal and ventral hippocampus and their functional interactions with the prefrontal cortex. Hippocampus 30:733–744.

Bolkvadze T, Pitkanen A (2012) Development of post-traumatic epilepsy after controlled cortical impact and lateral fluid-percussion-induced brain injury in the mouse. J Neurotrauma 29:789–812.

Buerge T, Weiss T (2004) Handling and Restraint. In: The Laboratory Mouse (Handbook of Experimental Animals) (Hedrich HJ, Bullock G, eds), pp 517–526: Elsevier.

Buzsaki G (1989) Two-stage model of memory trace formation: a role for “noisy” brain states. Neuroscience 31:551–570.

Cai DJ et al. (2016) A shared neural ensemble links distinct contextual memories encoded close in time. Nature 534:115–118.

Carter CS, Richardson A, Huffman DM, Austad S (2020) Bring Back the Rat! J Gerontol A Biol Sci Med Sci 75:405–415.

Chan KH, Morell JR, Jarrard LE, Davidson TL (2001) Reconsideration of the role of the hippocampus in learned inhibition. Behav Brain Res 119:111–130.

Chen LJ, Wang YJ, Chen JR, Tseng GF (2017) Hydrocephalus compacted cortex and hippocampus and altered their output neurons in association with spatial learning and memory deficits in rats. Brain Pathol 27:419–436.

Chen TW, Wardill TJ, Sun Y, Pulver SR, Renninger SL, Baohan A, Schreiter ER, Kerr RA, Orger MB, Jayaraman V, Looger LL, Svoboda K, Kim DS (2013) Ultrasensitive fluorescent proteins for imaging neuronal activity. Nature 499:295–300.

Chrobak JJ, Buzsaki G (1996) High-frequency oscillations in the output networks of the hippocampal-entorhinal axis of the freely behaving rat. J Neurosci 16:3056–3066.

Colgin LL (2013) Mechanisms and functions of theta rhythms. Annu Rev Neurosci 36:295–312.

Colgin LL, Kubota D, Jia Y, Rex CS, Lynch G (2004) Long-term potentiation is impaired in rat hippocampal slices that produce spontaneous sharp waves. J Physiol 558:953–961.

Corder G, Ahanonu B, Grewe BF, Wang D, Schnitzer MJ, Scherrer G (2019) An amygdalar neural ensemble that encodes the unpleasantness of pain. Science 363:276–281.

Cressant A, Besson M, Suarez S, Cormier A, Granon S (2007) Spatial learning in Long-Evans Hooded rats and C57BL/6J mice: different strategies for different performance. Behav Brain Res 177:22–29.

Csicsvari J, Hirase H, Czurko A, Mamiya A, Buzsaki G (1999) Fast network oscillations in the hippocampal CA1 region of the behaving rat. J Neurosci 19:RC20.

Dombeck DA, Harvey CD, Tian L, Looger LL, Tank DW (2010) Functional imaging of hippocampal place cells at cellular resolution during virtual navigation. Nat Neurosci 13:1433–1440.

Ego-Stengel V, Wilson MA (2007) Spatial selectivity and theta phase precession in CA1 interneurons. Hippocampus 17:161–174.

Eichenbaum H, Dudchenko P, Wood E, Shapiro M, Tanila H (1999) The hippocampus, memory, and place cells: is it spatial memory or a memory space? Neuron 23:209–226.

Ellenbroek B, Youn J (2016) Rodent models in neuroscience research: is it a rat race? Dis Model Mech 9:1079–1087.

Fenton AA, Kao H-Y, Neymotin SA, Olypher A, Vayntrub Y, Lytton WW, Ludvig N (2008) Unmasking the CA1 ensemble place code by exposures to small and large environments: more place cells and multiple, irregularly arranged, and expanded place fields in the larger space. Journal of Neuroscience 28:11250–11262.

Ferbinteanu J, Shapiro ML (2003) Prospective and retrospective memory coding in the hippocampus. Neuron 40:1227–1239.

Ferbinteanu J, Kennedy PJ, Shapiro ML (2006) Episodic memory—from brain to mind. Hippocampus 16:691–703.

Foster DJ, Knierim JJ (2012) Sequence learning and the role of the hippocampus in rodent navigation. Curr Opin Neurobiol 22:294–300.

Foster DJ, Morris RG, Dayan P (2000) A model of hippocampally dependent navigation, using the temporal difference learning rule. Hippocampus 10:1–16.

Franklin KBJ, Paxinos G (2013) Paxinos and Franklin’s The mouse brain in stereotaxic coordinates.

Frick KM, Stillner ET, Berger-Sweeney J (2000) Mice are not little rats: species differences in a one-day water maze task. Neuroreport 11:3461–3465.

Grienberger C, Konnerth A (2012a) Imaging calcium in neurons. Neuron 73:862–885.

Grienberger C, Konnerth A (2012b) Imaging calcium in neurons. Neuron 73:862–885.

Grosmark AD, Sparks FT, Davis MJ, Losonczy A (2021) Reactivation predicts the consolidation of unbiased long-term cognitive maps. Nat Neurosci 24:1574–1585.

Guo W, Zhang JJ, Newman JP, Wilson MA (2020) Latent learning drives sleep-dependent plasticity in distinct CA1 subpopulations. bioRxiv:2020.2002.2027.967794.

Guzowski JF, McNaughton BL, Barnes CA, Worley PF (1999) Environment-specific expression of the immediate-early gene Arc in hippocampal neuronal ensembles. Nat Neurosci 2:1120–1124.

Hainmueller T, Bartos M (2018) Parallel emergence of stable and dynamic memory engrams in the hippocampus. Nature 558:292–296.

Hamel EJ, Grewe BF, Parker JG, Schnitzer MJ (2015) Cellular level brain imaging in behaving mammals: an engineering approach. Neuron 86:140–159.

Hart EE, Blair GJ, O’Dell TJ, Blair HT, Izquierdo A (2020) Chemogenetic modulation and single-photon calcium imaging in anterior cingulate cortex reveal a mechanism for effort-based decisions. Journal of Neuroscience 40:5628–5643.

Hasselmo ME (2005) What is the function of hippocampal theta rhythm?--Linking behavioral data to phasic properties of field potential and unit recording data. Hippocampus 15:936–949.

Hasselmo ME, McClelland JL (1999) Neural models of memory. Curr Opin Neurobiol 9:184–188.

Hasselmo ME, Hay J, Ilyn M, Gorchetchnikov A (2002) Neuromodulation, theta rhythm and rat spatial navigation. Neural Netw 15:689–707.

Hirst WD, Abrahamsen B, Blaney FE, Calver AR, Aloj L, Price GW, Medhurst AD (2003) Differences in the central nervous system distribution and pharmacology of the mouse 5-hydroxytryptamine-6 receptor compared with rat and human receptors investigated by radioligand binding, site-directed mutagenesis, and molecular modeling. Molecular pharmacology 64:1295–1308.

Hok V, Poucet B, Duvelle É, Save É, Sargolini F (2016) Spatial cognition in mice and rats: similarities and differences in brain and behavior. Wiley Interdisciplinary Reviews: Cognitive Science 7:406–421.

Huang L, Ledochowitsch P, Knoblich U, Lecoq J, Murphy GJ, Reid RC, de Vries SE, Koch C, Zeng H, Buice MA (2021) Relationship between simultaneously recorded spiking activity and fluorescence signal in GCaMP6 transgenic mice. eLife 10:e51675.

Hunt RF, Scheff SW, Smith BN (2009) Posttraumatic epilepsy after controlled cortical impact injury in mice. Exp Neurol 215:243–252.

Hunt RF, Scheff SW, Smith BN (2010) Regionally localized recurrent excitation in the dentate gyrus of a cortical contusion model of posttraumatic epilepsy. J Neurophysiol 103:1490–1500.

Huxter J, Burgess N, O’Keefe J (2003) Independent rate and temporal coding in hippocampal pyramidal cells. Nature 425:828–832.

Iannaccone PM, Jacob HJ (2009) Rats! Dis Model Mech 2:206–210.

Ito R, Everitt BJ, Robbins TW (2005) The hippocampus and appetitive Pavlovian conditioning: effects of excitotoxic hippocampal lesions on conditioned locomotor activity and autoshaping. Hippocampus 15:713–721.

Jimenez JC, Berry JE, Lim SC, Ong SK, Kheirbek MA, Hen R (2020) Contextual fear memory retrieval by correlated ensembles of ventral CA1 neurons. Nature communications 11:3492.

Josselyn SA, Tonegawa S (2020) Memory engrams: Recalling the past and imagining the future. Science 367.

Jun H, Bramian A, Soma S, Saito T, Saido TC, Igarashi KM (2020) Disrupted Place Cell Remapping and Impaired Grid Cells in a Knockin Model of Alzheimer’s Disease. Neuron 107:1095–1112 e1096.

Jupp B, Caprioli D, Dalley JW (2013) Highly impulsive rats: modelling an endophenotype to determine the neurobiological, genetic and environmental mechanisms of addiction. Dis Model Mech 6:302–311.

Kinsky NR, Sullivan DW, Mau W, Hasselmo ME, Eichenbaum HB (2018) Hippocampal Place Fields Maintain a Coherent and Flexible Map across Long Timescales. Current biology : CB 28:3578–3588 e3576.

Kinsky NR, Mau W, Sullivan DW, Levy SJ, Ruesch EA, Hasselmo ME (2020) Trajectory-modulated hippocampal neurons persist throughout memory-guided navigation. Nature communications 11:2443.

Klaus A, Martins GJ, Paixao VB, Zhou P, Paninski L, Costa RM (2017) The Spatiotemporal Organization of the Striatum Encodes Action Space. Neuron 95:1171–1180 e1177.

Kudrimoti HS, Barnes CA, McNaughton BL (1999) Reactivation of hippocampal cell assemblies: effects of behavioral state, experience, and EEG dynamics. J Neurosci 19:4090–4101.

Leutgeb S, Leutgeb JK, Treves A, Moser MB, Moser EI (2004) Distinct ensemble codes in hippocampal areas CA3 and CA1. Science 305:1295–1298.

Levy SJ, Kinsky NR, Mau W, Sullivan DW, Hasselmo ME (2021) Hippocampal spatial memory representations in mice are heterogeneously stable. Hippocampus 31:244–260.

Lopes G, Bonacchi N, Frazao J, Neto JP, Atallah BV, Soares S, Moreira L, Matias S, Itskov PM, Correia PA, Medina RE, Calcaterra L, Dreosti E, Paton JJ, Kampff AR (2015) Bonsai: an event-based framework for processing and controlling data streams. Front Neuroinform 9:7.

Maier N, Nimmrich V, Draguhn A (2003) Cellular and network mechanisms underlying spontaneous sharp wave-ripple complexes in mouse hippocampal slices. J Physiol 550:873–887.

Malvache A, Reichinnek S, Villette V, Haimerl C, Cossart R (2016) Awake hippocampal reactivations project onto orthogonal neuronal assemblies. Science 353:1280–1283.

Markus EJ, Barnes CA, McNaughton BL, Gladden VL, Skaggs WE (1994) Spatial information content and reliability of hippocampal CA1 neurons: effects of visual input. Hippocampus 4:410–421.

Mau W, Sullivan DW, Kinsky NR, Hasselmo ME, Howard MW, Eichenbaum H (2018) The Same Hippocampal CA1 Population Simultaneously Codes Temporal Information over Multiple Timescales. Current biology : CB 28:1499–1508 e1494.

Maurer AP, Cowen SL, Burke SN, Barnes CA, McNaughton BL (2006) Organization of hippocampal cell assemblies based on theta phase precession. Hippocampus 16:785–794.

McEchron MD, Disterhoft JF (1999) Hippocampal encoding of non-spatial trace conditioning. Hippocampus 9:385–396.

McNaughton BL, Barnes CA, O’Keefe J (1983) The contributions of position, direction, and velocity to single unit activity in the hippocampus of freely-moving rats. Experimental brain research 52:41–49.

McNaughton BL, Battaglia FP, Jensen O, Moser EI, Moser MB (2006) Path integration and the neural basis of the ’cognitive map’. Nat Rev Neurosci 7:663–678.

Meshulam L, Gauthier JL, Brody CD, Tank DW, Bialek W (2017) Collective Behavior of Place and Non-place Neurons in the Hippocampal Network. Neuron 96:1178–1191 e1174.

Moser EI, Moser M-B, McNaughton BL (2017) Spatial representation in the hippocampal formation: a history. Nature neuroscience 20:1448–1464.

Mou X, Cheng J, Yu YS, Kee SE, Ji D (2018) Comparing mouse and rat hippocampal place cell activities and firing sequences in the same environments. Frontiers in cellular neuroscience 12:332.

Moyer JR, Jr., Deyo RA, Disterhoft JF (1990) Hippocampectomy disrupts trace eye-blink conditioning in rabbits. Behav Neurosci 104:243–252.

Nathanson JL, Yanagawa Y, Obata K, Callaway EM (2009) Preferential labeling of inhibitory and excitatory cortical neurons by endogenous tropism of adeno-associated virus and lentivirus vectors. Neuroscience 161:441–450.

O’Keefe J (1976) Place units in the hippocampus of the freely moving rat. Exp Neurol 51:78–109.

Olypher AV, Lansky P, Muller RU, Fenton AA (2003) Quantifying location-specific information in the discharge of rat hippocampal place cells. J Neurosci Methods 127:123–135.

Parker CC, Chen H, Flagel SB, Geurts AM, Richards JB, Robinson TE, Woods LCS, Palmer AA (2014) Rats are the smart choice: rationale for a renewed focus on rats in behavioral genetics. Neuropharmacology 76:250–258.

Podor B, Hu Y-l, Ohkura M, Nakai J, Croll R, Fine A (2015a) Comparison of genetically encoded calcium indicators for monitoring action potentials in mammalian brain by two-photon excitation fluorescence microscopy. Neurophotonics 2:021014.

Podor B, Hu YL, Ohkura M, Nakai J, Croll R, Fine A (2015b) Comparison of genetically encoded calcium indicators for monitoring action potentials in mammalian brain by two-photon excitation fluorescence microscopy. Neurophotonics 2:021014.

Pritchard A, Hornung R, Kramer PR (2021) Measuring calcium activity within individual neurons within the rat thalamus. MethodsX 8:101273.

Resendez SL, Jennings JH, Ung RL, Namboodiri VM, Zhou ZC, Otis JM, Nomura H, McHenry JA, Kosyk O, Stuber GD (2016) Visualization of cortical, subcortical and deep brain neural circuit dynamics during naturalistic mammalian behavior with head-mounted microscopes and chronically implanted lenses. Nature protocols 11:566–597.

Rich PD, Liaw HP, Lee AK (2014) Place cells. Large environments reveal the statistical structure governing hippocampal representations. Science 345:814–817.

Rickgauer JP, Deisseroth K, Tank DW (2014) Simultaneous cellular-resolution optical perturbation and imaging of place cell firing fields. Nat Neurosci 17:1816–1824.

Rosenfeld CS, Ferguson SA (2014) Barnes maze testing strategies with small and large rodent models. J Vis Exp:e51194.

Rosenzweig ES, Redish AD, McNaughton BL, Barnes CA (2003) Hippocampal map realignment and spatial learning. Nature neuroscience 6:609–615.

Routh BN, Johnston D, Harris K, Chitwood RA (2009) Anatomical and electrophysiological comparison of CA1 pyramidal neurons of the rat and mouse. Journal of neurophysiology 102:2288–2302.

Schimanski LA, Lipa P, Barnes CA (2013) Tracking the course of hippocampal representations during learning: when is the map required? J Neurosci 33:3094–3106.

Schoenfeld G, Carta S, Rupprecht P, Ayaz A, Helmchen F (2021) In vivo calcium imaging of CA3 pyramidal neuron populations in adult mouse hippocampus. eNeuro.

Scott BB, Thiberge SY, Guo C, Tervo DGR, Brody CD, Karpova AY, Tank DW (2018) Imaging cortical dynamics in GCaMP transgenic rats with a head-mounted widefield macroscope. Neuron 100:1045–1058. e1045.

Scoville WB, Milner B (1957) Loss of recent memory after bilateral hippocampal lesions. Journal of neurology, neurosurgery, and psychiatry 20:11–21.

Sheintuch L, Geva N, Baumer H, Rechavi Y, Rubin A, Ziv Y (2020) Multiple Maps of the Same Spatial Context Can Stably Coexist in the Mouse Hippocampus. Current biology : CB 30:1467–1476 e1466.

Sheintuch L, Rubin A, Brande-Eilat N, Geva N, Sadeh N, Pinchasof O, Ziv Y (2017) Tracking the Same Neurons across Multiple Days in Ca(2+) Imaging Data. Cell reports 21:1102–1115.

Shemesh OA et al. (2020) Precision Calcium Imaging of Dense Neural Populations via a Cell-Body-Targeted Calcium Indicator. Neuron 107:470–486 e411.

Shim I, Ha Y, Chung JY, Lee HJ, Yang KH, Chang JW (2003) Association of learning and memory impairments with changes in the septohippocampal cholinergic system in rats with kaolin-induced hydrocephalus. Neurosurgery 53:416–425; discussion 425.

Shin JH, Song M, Paik SB, Jung MW (2020) Spatial organization of functional clusters representing reward and movement information in the striatal direct and indirect pathways. Proc Natl Acad Sci U S A 117:27004–27015.

Shirvalkar PR, Rapp PR, Shapiro ML (2010) Bidirectional changes to hippocampal theta–gamma comodulation predict memory for recent spatial episodes. Proceedings of the National Academy of Sciences 107:7054–7059.

Shuman T et al. (2020) Breakdown of spatial coding and interneuron synchronization in epileptic mice. Nat Neurosci 23:229–238.

Silva AJ (2017) Miniaturized two-photon microscope: seeing clearer and deeper into the brain. Light: Science & Applications 6:e17104–e17104.

Smith DM, Mizumori SJ (2006) Hippocampal place cells, context, and episodic memory. Hippocampus 16:716–729.

Snyder JS, Choe JS, Clifford MA, Jeurling SI, Hurley P, Brown A, Kamhi JF, Cameron HA (2009) Adult-born hippocampal neurons are more numerous, faster maturing, and more involved in behavior in rats than in mice. Journal of Neuroscience 29:14484–14495.

Souza BC, Pavão R, Belchior H, Tort AB (2018) On information metrics for spatial coding. Neuroscience 375:62–73.

Squire LR (1992) Memory and the hippocampus: a synthesis from findings with rats, monkeys, and humans. Psychol Rev 99:195–231.

Stefanini F, Kushnir L, Jimenez JC, Jennings JH, Woods NI, Stuber GD, Kheirbek MA, Hen R, Fusi S (2020) A Distributed Neural Code in the Dentate Gyrus and in CA1. Neuron 107:703–716 e704.

Steinmetz NA et al. (2017) Aberrant Cortical Activity in Multiple GCaMP6-Expressing Transgenic Mouse Lines. eNeuro 4.

Swanson L (2004) Brain maps: structure of the rat brain: Gulf Professional Publishing.

Talbot ZN, Sparks FT, Dvorak D, Curran BM, Alarcon JM, Fenton AA (2018) Normal CA1 Place Fields but Discoordinated Network Discharge in a Fmr1-Null Mouse Model of Fragile X Syndrome. Neuron 97:684–697 e684.

Thevenaz P, Ruttimann UE, Unser M (1998) A pyramid approach to subpixel registration based on intensity. IEEE Trans Image Process 7:27–41.

Tian L, Hires SA, Mao T, Huber D, Chiappe ME, Chalasani SH, Petreanu L, Akerboom J, McKinney SA, Schreiter ER (2009a) Imaging neural activity in worms, flies and mice with improved GCaMP calcium indicators. Nature methods 6:875–881.

Tian L, Hires SA, Mao T, Huber D, Chiappe ME, Chalasani SH, Petreanu L, Akerboom J, McKinney SA, Schreiter ER, Bargmann CI, Jayaraman V, Svoboda K, Looger LL (2009b) Imaging neural activity in worms, flies and mice with improved GCaMP calcium indicators. Nat Methods 6:875–881.

Vann SD (2013) Dismantling the Papez circuit for memory in rats. eLife 2:e00736.

Vengeliene V, Bilbao A, Spanagel R (2014) The alcohol deprivation effect model for studying relapse behavior: a comparison between rats and mice. Alcohol 48:313–320.

Voigts J, Siegle JH, Pritchett DL, Moore CI (2013) The flexDrive: an ultra-light implant for optical control and highly parallel chronic recording of neuronal ensembles in freely moving mice. Front Syst Neurosci 7:8.

Voigts J, Newman JP, Wilson MA, Harnett MT (2020) An easy-to-assemble, robust, and lightweight drive implant for chronic tetrode recordings in freely moving animals. J Neural Eng 17:026044.

Wang X, Liu Y, Li X, Zhang Z, Yang H, Zhang Y, Williams PR, Alwahab NSA, Kapur K, Yu B, Zhang Y, Chen M, Ding H, Gerfen CR, Wang KH, He Z (2017) Deconstruction of Corticospinal Circuits for Goal-Directed Motor Skills. Cell 171:440–455 e414.

Whishaw IQ (1995) A comparison of rats and mice in a swimming pool place task and matching to place task: some surprising differences. Physiol Behav 58:687–693.

Whishaw IQ, Metz GA, Kolb B, Pellis SM (2001) Accelerated nervous system development contributes to behavioral efficiency in the laboratory mouse: a behavioral review and theoretical proposal. Developmental Psychobiology: The Journal of the International Society for Developmental Psychobiology 39:151–170.

Wikenheiser AM, Redish AD (2015) Hippocampal theta sequences reflect current goals. Nat Neurosci 18:289–294.

Wilent WB, Nitz DA (2007) Discrete place fields of hippocampal formation interneurons. J Neurophysiol 97:4152–4161.

Wilson MA, McNaughton BL (1993) Dynamics of the hippocampal ensemble code for space. Science 261:1055–1058.

Wirtshafter HS, Wilson MA (2019) Locomotor and Hippocampal Processing Converge in the Lateral Septum. Current biology : CB 29:3177–3192.

Wirtshafter HS, Wilson MA (2020) Differences in reward biased spatial representations in the lateral septum and hippocampus. eLife 9.

Wirtshafter HS, Wilson MA (2021) Lateral septum as a nexus for mood, motivation, and movement. Neurosci Biobehav Rev 126:544–559.

Witharana WK, Cardiff J, Chawla MK, Xie JY, Alme CB, Eckert M, Lapointe V, Demchuk A, Maurer AP, Trivedi V, Sutherland RJ, Guzowski JF, Barnes CA, McNaughton BL (2016) Nonuniform allocation of hippocampal neurons to place fields across all hippocampal subfields. Hippocampus 26:1328–1344.

Yang Y, Liu N, He Y, Liu Y, Ge L, Zou L, Song S, Xiong W, Liu X (2018) Improved calcium sensor GCaMP-X overcomes the calcium channel perturbations induced by the calmodulin in GCaMP. Nature communications 9:1504.

Ylinen A, Bragin A, Nadasdy Z, Jando G, Szabo I, Sik A, Buzsaki G (1995) Sharp wave-associated high-frequency oscillation (200 Hz) in the intact hippocampus: network and intracellular mechanisms. J Neurosci 15:30–46.

Zhang K, Ginzburg I, McNaughton BL, Sejnowski TJ (1998) Interpreting neuronal population activity by reconstruction: unified framework with application to hippocampal place cells. J Neurophysiol 79:1017–1044.

Zhang X, Li B (2018) Population coding of valence in the basolateral amygdala. Nature communications 9:5195.

Zhou H, Neville KR, Goldstein N, Kabu S, Kausar N, Ye R, Nguyen TT, Gelwan N, Hyman BT, Gomperts SN (2019) Cholinergic modulation of hippocampal calcium activity across the sleep-wake cycle. eLife 8.

Zhou P, Resendez SL, Rodriguez-Romaguera J, Jimenez JC, Neufeld SQ, Giovannucci A, Friedrich J, Pnevmatikakis EA, Stuber GD, Hen R, Kheirbek MA, Sabatini BL, Kass RE, Paninski L (2018) Efficient and accurate extraction of in vivo calcium signals from microendoscopic video data. eLife 7.

Ziv Y, Burns LD, Cocker ED, Hamel EO, Ghosh KK, Kitch LJ, El Gamal A, Schnitzer MJ (2013) Long-term dynamics of CA1 hippocampal place codes. Nat Neurosci 16:264–266.

